# Phylogenetically diverse introgression drives subtle population structure in Pacific rockfishes

**DOI:** 10.1101/2025.09.23.678143

**Authors:** Nathan T. B. Sykes, R. Nicolas Lou, Matthew R. Siegle, Peter H. Sudmant, Wesley A. Larson, Gregory L. Owens

## Abstract

Genomic methods have shown that admixture and introgression is common across animal taxa. Pacific rockfishes, genus *Sebastes*, are group of commercially important species that primarily inhabit inshore, shelf, and slope habitats along the North American west coast. Among these, Copper and Quillback Rockfishes (abbreviated to Copper and Quillback) are closely related species known to hybridize, particularly within the Salish Sea in North America’s Pacific Northwest. Here, we investigate genetic population structure and introgression patterns in Copper and Quillback from Alaska to California. Using whole-genome resequencing (WGS) across a broad geographic range, we seek to (1) compare population structure between these species, and (2) assess how introgression affects population structure patterns. Our analyses reveal that Copper exhibit much higher levels of population differentiation compared to Quillback, especially separating Salish Sea samples from all other populations. In contrast, Quillback populations appear to be nearly panmictic, with lower overall differentiation. Surprisingly, we detected signatures of introgression from 13 other rockfish species in Copper and 16 species in Quillback. This introgression was highly regional suggesting hybridization depended on geographic context and congener ranges. Yelloweye Rockfish introgression drives the strongest signal of regional population structure in Quillback. These findings provide novel insights into the range-wide genetic structure of these species and highlight that hybridization in *Sebastes* is phylogenetically broader than previously appreciated.

## Introduction

Marine fishes tend toward subtle population structure, even panmixia. This is due in part to their large population sizes and high dispersal capacity, as well as the relative continuity of marine habitat, reducing barriers to gene flow (Nielsen et al., 2009; Waples, 1998). Some species may exhibit population differentiation reflecting their unique life history, fragmented habitat, or capacity for dispersal. Anadromous Pacific salmon, for example, have population structure reflecting their spawning site fidelity (Xuereb et al., 2022), run timing (Narum et al., 2023), or life cycle (Tarpey et al., 2018). In the absence of strong reproductive isolation based on geography, though, intraspecific differentiation is harder to detect. High rates of migration and corresponding high gene flow can lead to largely homogenized populations, both at subcontinental and oceanic scales (Díaz-Arce et al., 2024; Natasha et al., 2022; Tovar Verba et al., 2022). This presents a challenge for fisheries managers, who seek insight into patterns of dispersal to establish conservation units. Genetic analyses, such as those which leverage sequence divergence within allozymes or microsatellites, have historically been applied to answer these questions, but are limited to only a handful of genetic markers (Buonaccorsi et al., 2004; Gilbert-Horvath et al., 2006; Gao et al., 2016). However, subtle population structure may be challenging to identify with small numbers of markers. Genomic techniques provide researchers with improved resolution to study population structure in marine fishes (Bernatchez et al., 2017; Longo et al., 2020), but many species have not been analyzed with these techniques, leaving fisheries managers to rely on decades-old genetic analyses in many cases. Given the increasing effectiveness and affordability of genomic resources, it is essential to close this knowledge gap, particularly for overfished and threatened species.

North Eastern Pacific rockfishes, genus *Sebastes*, comprise many economically, socially, and culturally important species, which inhabit inshore, shelf, and slope habitats along the North American West coast (Love et al., 2002, Hyde & Vetter, 2007). Among inshore rockfishes, Copper (*S. caurinus*), and Quillback (*S. maliger*) rockfishes (abbreviated to Copper or Quillback) are two closely related species that hybridize extensively, particularly within the Salish Sea (Seeb, 1998; Schwenke et al., 2018; Wray et al., 2024). Both species are broadly distributed from Alaska to California, although Copper exhibit a more southerly distribution extending into Baja California.

In Canada and the United States, rockfish stocks are often managed along political boundaries (between countries and states) and at regional scales whereby fish in the Salish Sea, for example, are considered a separate stock from those of the greater coast for many species. While for Copper and Quillback, the latter delineation was originally made to reflect exploitation history and not genomic differentiation between the regions, earlier work suggests that the management units reflect some degree of reproductive isolation (Buonaccorsi et al., 2002, 2005; Seeb, 1998). The trend is similar in Yelloweye Rockfish, which in the Salish Sea are genetically distinct from the outer coast (Siegle et al., 2013; Andrews et al., 2018). Retention of Copper and Quillback in the Salish Sea is prohibited in US waters and highly limited in Canadian waters. Copper in outside waters from Alaska to California are managed based on the following spatial boundaries: Alaska, British Columbia, Oregon and Washington, and California (NMFS, 2023). Quillback in outside waters from Alaska to California are managed based on the following spatial boundaries: Alaska, British Columbia, Washington, Oregon, and California. Quillback were recently designated as overfished in California waters, prompting fisheries closures.

Pacific rockfishes exhibit characteristics that can result in spatially discrete population structure. Small adult home ranges, courtship rituals, internal fertilization, parturition of live offspring, oceanographic effects on larval dispersal, and nearshore early life-history stages may all reasonably conspire to limit gene flow over large distances and promote genetic differentiation (Buonaccorsi et al., 2002, 2005; Hannah & Rankin, 2011; Helmstetter et al., 2016; Love et al., 2002; Matthews, 1990; Tolimieri et al., 2009). However, rockfish can also display very low genetic differentiation across large spatial scales, likely due to their relatively long pelagic larval dispersal (lasting up to several months) that can facilitate gene flow over significant distances (Siegle et al., 2013). Previous examinations of population structure in Copper and Quillback have been technologically or regionally limited. Allozyme and microsatellite based genetic analyses suggest that both species exhibit significant population structure only when Puget Sound is considered (Buonaccorsi et al., 2002, 2005; Seeb, 1998). Another study that examined differentiation at microsatellite loci in Copper found replicate divergence between pairs of inlet and coast samples on the west coast of Vancouver Island (Dick et al., 2014). Most recently, a genomic study focusing on the Salish Sea and Puget Sound identified two distinct populations in Quillback, one largely in southern Puget Sound and another in the Salish Sea (Wray et al., 2024). They also uncovered three discrete populations in Copper, again largely separating Puget Sound from the Salish Sea. Together, this could be taken as an indication that Puget Sound and the Salish Sea harbour unique genetic diversity, but it has not been put in the context of the rest of the species range.

Despite the relatively recent radiation of rockfish species, hybridization is thought to be relatively rare in the genus with two exceptions: Puget Sound and recent species pairs (Love et al., 2002). In Puget Sound, hybrid fish are more common (Seeb, 1998). This has been mostly studied between the Quillback, Copper and Brown Rockfish and is thought to be caused by a combination of anoxic conditions, reduced habitat availability and/or differences in population size between species (Seeb, 1998; Buonaccorsi et al., 2002; Buonaccorsi et al., 2005; Wray et al., 2024; Schwenke et al., 2018). Hybridization has also been noted between closely related species pairs, including between *S. chrysomelas* and *S. carnatus* (Buonaccorsi et al., 2005), *S. vulpes* and *S. zonatus* (Muto et al., 2013), *S. mentella* and *S. viviparous* (Artamonova et al., 2013). The extent to which hybridization has played a role in speciation or adaptation in the genus is currently an open question.

This study leverages whole genome resequencing (WGS) of Copper and Quillback to describe their genetic diversity and population structure throughout their native range. Through this international sampling effort, we seek to: (1) compare population structure between each species, (2) quantify introgression and its effect on population structure. Through the first range-wide characterization of genetic diversity in these species, we provide novel insight into their patterns of dispersal, differentiation, and introgression.

## Methods

### Sample collection

Samples were collected by various means, including in annual longline surveys conducted by Fisheries and Oceans Canada (DFO) and the US National Oceanic and Atmospheric Administration (NOAA), and in individual rod-and-reel catch efforts conducted by authors and collaborators (Anderson et al., 2019). In total, 113 Copper (*S. caurinus*) and 200 Quillback (*S. maliger*) rockfish were collected between June 1994 and August 2022. Samples were sorted into eight regional bins: Alaska (AK), Hecate Straight (HS), Queen Charlotte Sound (QCS), Western Vancouver Island (WVI), Salish Sea (SS), Puget Sound (PS), Washington and Oregon (WA/OR), and California (CA; Fig. 1B). Throughout the paper, we refer to samples in Puget Sound and Salish Sea as ‘inside’ and all others as ‘outside’. Additionally, as outgroups we included five Gopher (*S. carnatus*) and five Black-and-yellow (*S. chrysomelas*) Rockfish samples collected from the same sources which, while closely related, only range from southern Oregon to Baja California (Hyde and Vetter, 2007).

**Figure 1:**
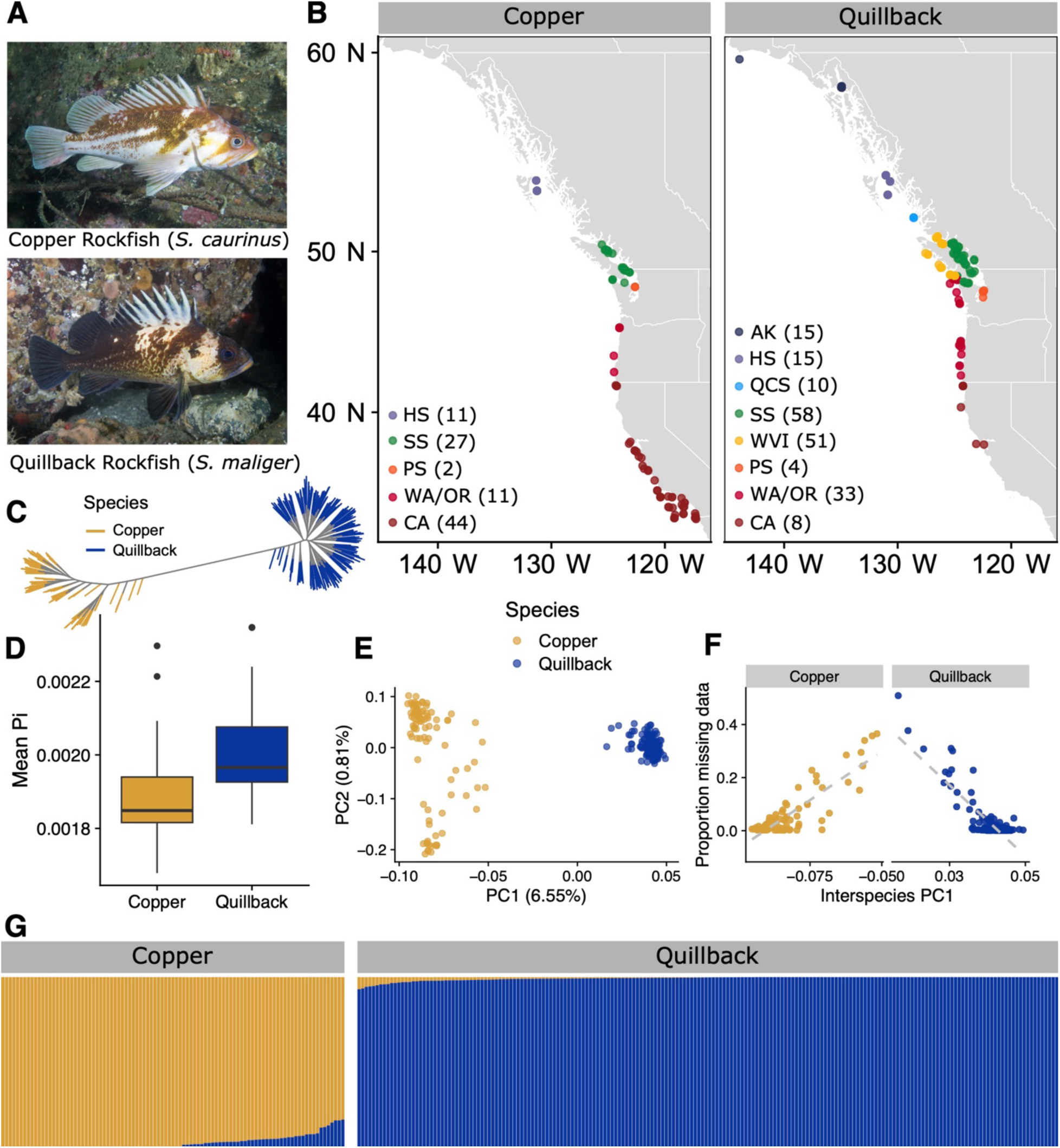
Sampling and interspecific differentiation. **A)** Underwater photographs of each species by Andy Murch at Big Fish Expeditions. **B)** Sampling distribution and geographic bins. The number in parentheses represents the samples retained after filtering from each population: Alaska (AK), Hecate Strait (HS), Queen Charlotte Sound (QCS), Salish Sea (SS), Western Vancouver Island (WVI), Puget Sound (PS), Washington and Oregon (WA/OR) and California (CA).**C)** Unrooted consensus tree of all individuals retained in the data set, coloured by species. **D)** Mean nucleotide diversity (π) of chromosomes in both species. **E)** Principal component analysis of all samples, coloured by species. **F)** Intermediate PC1 score is caused by increased missing data. **G)** Combined ADMIXTURE analysis of all samples for K = 2.

### Library prep and sequencing

All whole genome resequencing library preparation was performed using methods similar to Baym et al. (2015), Therkildsen & Palumbi (2017), and Euclide et al., (2023). Genomic DNA input was normalized to 10 ng per individual. Per Euclide et al. (2023), sample purification, product normalization and pooling were conducted with SequalPrep plates (ThermoFisher Scientific: Waltham, MA, USA) rather than AMPure XP beads (Beckman Coulter: Brea, CA, USA). Prior to sequencing, the final pooled library was visualized on a 2% agarose E-gel (ThermoFisher Scientific) and quantified with the Qubit HS dsDNA Assay Kit (ThermoFisher Scientific).

Sequencing of an initial batch of 113 Copper and 152 Quillback, as well as the Gopher and Black-and-yellow Rockfish samples, occurred at Novogene (Sacramento, CA, USA) on Illumina NovaSeq S4. An additional 48 Quillback samples were also sequenced on a Novaseq with S4 chemistry by the University of Oregon’s Genomic & Cell Characterization Core Facility (GC3F; Eugene, OR, USA). Samples were sequenced on lanes containing 96 rockfish each and were multiplexed with samples for other projects.

### Alignment, variant calling and quality control

Alignment and initial quality control were done using the Snakemake pipeline *grenepipe* v0.10.0 (Czech & Exposito-Alonso, 2021). All tools were run using default settings unless specified otherwise. First, adapter sequences were trimmed using *fastp* v0.20.0 (Chen, 2023). Reads were then aligned to a Korean rockfish (*S. schlegelii*) genome acquired from the Chinese National GeneBank Database (He et al., 2019; CNA0000824) using *bwa-mem* v2.2.1 (Vasimuddin et al., 2019). This reference genome was chosen since it is an outgroup of all sequenced species and prevents reference bias in interspecies comparisons. Overlapping read pairs were clipped using *BamUtil clipOverlap* v1.0.15 (Jun et al., 2015), and duplicated reads were removed using *picard MarkDuplicates* v2.23.0 (Broad Institute, 2019) with the parameters *-Xmx80g REMOVE_DUPLICATES=true, VALIDATION_STRINGENCY=SILENT. FastQC* v0.11.9 (S. Andrews, 2010), *Qualimap* v2.2.2a (Okonechnikov et al., 2016), *picard CollectMultipleMetrics* v2.23.0 (Broad Institute, 2019), and *MultiQC* v1.10.1 (Ewels et al., 2016) were used to assess the quality of the data throughout the pipeline.

Variants were called using *freebayes* v1.3.6 (Garrison & Marth, 2012), for which we skipped sites with overall read depth in excess of 20x the total number of samples since average depth was 6.04x, and set both minimum mapping and base quality to 20 (*-g 6260 -m 20 -q 20).* Monomorphic sites were retained in the unfiltered dataset for estimates of genetic diversity. We then used *vcftools* v0.1.16 (Danecek et al., 2011) to extract individual and site-based statistics for the remaining samples and excluded some poor-quality samples (>25% missing data). In total, 18 Copper and six Quillback were excluded from analysis due to missing data or suspected sample contamination. The final total number of samples retained were 95 Copper and 194 Quillback (Fig. 1B; Table S1), as well as five samples each of Gopher and Black-and-yellow.

We then filtered the data for downstream analyses using both *bcftools* v1.20 (Danecek et al., 2021) and *vcftools* (Danecek et al., 2011); we retained only biallelic SNPs with a minor allele frequency greater than 5%, with maximum 25% missing data, minimum genotype read depth of five, maximum genotype read depth of 50, and QUAL ≥ 20. This filtering regime was applied to the whole dataset for interspecific analyses, as well as to each species separately for intraspecific analyses. After filtering, 1,047,615 SNPs were retained in the interspecific dataset, while 350,889 SNPs were retained in Copper and 1,447,179 SNPs in Quillback. The above computations and most downstream analyses were parallelized using *GNU parallel* (Tange, 2023).

### Population structure, differentiation, and genetic diversity

For both inter- and intraspecific population structure analyses, we performed principal component analyses (PCAs) in *R* v4.2.2 (R Core Team) with the *SNPRelate* package v3.13 (Zheng et al., 2012), and assessed ancestry proportions of our samples with *ADMIXTURE* v1.3.0 (Alexander et al., 2009). For ADMIXTURE, sites in high linkage disequilibrium were pruned using *PLINK* v1.90 (Purcell et al., 2007) in 10 kbp windows, with a maximum r-squared of 0.1 between retained sites in order to reduce redundancy and avoid overrepresentation of linked regions. For PCA, sites were pruned for linkage using snpgdsLDpruning in SNPRelate with an r-squared threshold of 0.2 within 500 kbp. For the interspecific PCA, we additionally filtered SNPs to require a maximum of 5% missing data because the signal of missing data conflated with introgression.

To generate a phylogeny of all samples in the dataset, we thinned and converted the full filtered dataset using a custom *perl* v5.32.1 (Wall et al., 2020) script, which retained 5% of sites. We then used iqtree2 v2.3.6 (Minh et al., 2020) with 1000 *Ultrafast* bootstraps (Minh et al., 2013; 2020) to generate a consensus tree (-m “GTR+ASC” -st DNA -B 1000).

Next, we sought to characterize the genomic differentiation both between and within the study species. To achieve this, we used a custom *perl* script to calculate F_ST_ for each position in the genome and summarized the results on a per-chromosome basis in *R* (Samuk et al., 2017). This analysis was applied to both species together and separately. Intraspecific F_ST_ was first calculated between geographic population bins. Then, for more granular comparisons, sampling coordinates were rounded to the nearest degree to make bins that included at least two samples. Eight Quillback and five Copper geographic bins included only a single sample so were excluded. To test for isolation by distance, we plotted F_ST_ versus Haversine distance for all coordinates. Since we found a major genetic barrier between the inside and outside waters, we grouped comparisons by inside-inside, inside-outside and outside-outside. To test for significance, we used a Mantel test and partial Mantel test, including inside/outside comparison type as a covariate using the vegan package in R (Sokal 1979; Oksanen et al. 2013). We observed that Quillback had significantly lower population differentiation than Copper, so we used *pixy* v1.2.7 (Korunes & Samuk, 2021) to estimate nucleotide diversity (π) in both species.

Given that differentiation was highest in comparisons made between fish in the Salish Sea (including Puget Sound) and outside of it, another pairwise F_ST_ calculation was conducted between all Salish Sea and Puget Sound samples and all others. We also subset samples inside and outside of the Salish Sea and Puget Sound for both species to run PCA and ADMIXTURE and test for regional structure.

### Reconciling introgression and population structure

Previous studies suggested that population structure in rockfish can be caused by introgression (Wray et al., 2024), so we conducted formal genome-wide tests of introgression using Patterson’s D, also known as the ABBA-BABA test, in *Dsuite* (Malinsky et al., 2021). This test selects two closely related species or populations (P1 and P2) and asks if one shares more derived alleles with a third species (P3). Without gene flow, both P1 and P2 should share equal numbers of derived alleles with P3 and D will be zero. If there is gene flow between P2 and P3, then D will be positive; conversely gene flow between P1 and P3 will result in a negative D. Importantly, if P1 or P2 has received gene flow from another source, this can also affect D (Tricou et al., 2022).

For our initial test, we assigned individuals to one of four groups based on the interspecific ADMIXTURE analysis; pure types from each species represented two of the groups, while any degree of mixed ancestry from ADMIXTURE would result in assignment to an admixed group for each species. We then used the *Dtrios* command to test whether samples identified as admixed were closer to their congener than non-admixed samples. The Korean rockfish reference genome was included as a sample in the vcf and specified as the outgroup.

Surprisingly, we found that the Quillback samples identified as admixed using ADMIXTURE shared fewer derived alleles with Copper than non-admixed samples. To explore this result and reconcile population structure and introgression, we conducted individual ABBA-BABA tests on all samples with the following sample order. For this, each individual sample was placed in the P2 position, and all other samples of the same species were put into the P1 position. All samples of the other species (Quillback or Copper) were placed in the P3 position (Figure 3C,D). The resulting scores represent a relative amount of allele sharing, compared to all other samples. Importantly, these scores do not reflect the absolute amount of introgression. For example, if all samples have 10% admixed ancestry, then all individual D-scores would be zero because they do not differ in admixture.

While the ABBA-BABA test is generally not direction (i.e. P2 into P3 or P3 into P2 are interchangeable), we believe our testing setup largely captures recent P3 into P2 introgression. For example, if a single sample is admixed, when placed in the P2 position it will share derived alleles with P3 that are not found in the other samples (placed in P1). If P3 is admixed with P1/P2 species, then it will largely have derived alleles found in both P1 and P2. There would only be a significant test if P1 and P2 separated different subpopulations of the species and the source of the P3 admixture was one subpopulation. In our tested species, there is very little population structure, so this scenario is unlikely.

### Genus-wide search for introgression donors

Our D-statistic suggested that Quillback contained introgression from a source other than Copper. To explore this, we selected publicly available whole-genome sequencing data for 51 species of Northeast Pacific Rockfish from across the genus (Table S2). Each sample was downsampled to 240 million paired-end reads if it had greater read depth to reduce processing time. Samples were processed as before, but SNP-calling was done in two datasets. Quillback samples plus genus-wide set, and Copper samples plus genus-wide set. In each dataset, a single high-coverage Copper or Quillback was added. After variant calling, we retained only biallelic SNPs with a minor allele frequency greater than 2%, with maximum 20% missing data, minimum read depth of five, maximum read depth of 60, and QUAL ≥ 20. We then repeated our individual D-stat analysis, isolating each Quillback/Copper sample and testing it against all others. We focused on comparisons in the following arrangement: P1: All other Quillback/Copper samples, P2: Target Quillback/Copper sample, P3: other *Sebastes* species. See the results section for a detailed interpretation of the test results.

## Results

### Interspecific analyses

Sequencing yielded an average read depth per sample of 5.25 in Copper, 6.83 in Quillback. All sequence data is available on the SRA (Table S1). In the interspecific PCA, the first principal component (PC1) separates samples along species lines and explains 6.55% of the total variance (percent variance explained; PVE). PC2 (0.8% PVE), separates Copper into their Salish Sea and coastal populations (Fig. 1E). Although some samples of both species drifted towards the center of the PCA, this was driven by missing data (Fig 1F). We found slightly higher diversity in Quillback (π = 0.002) over Copper (π = 0.00188) (Fig. 1D). An unrooted consensus tree of the full sample set reveals a pattern consistent with the PCA, whereby Quillback nodes radiate more from a central point than do the more differentiated Copper (Fig. 1C). Interspecific ADMIXTURE results indicate that there are 110 Quillback with some proportion of Copper ancestry (none exceeding 7.2%), and that there are 45 admixed Copper with up to 15.7% Quillback ancestry (Fig. 1E), although these results were not fully corroborated by further analyses. At K=3, the third group largely separated inside vs outside Copper (Fig. S1). At K=4 and 5, Quillback samples are appear to be largely admixed between three groups with little geographic patterning. Overall interspecific F_ST_ was calculated to be 0.333.

### Intraspecific population structure and differentiation

Intraspecific principal component analyses reveal that Copper have more defined population structure than Quillback (Fig. 2A). The first principal component in Copper (3.83% PVE) divides fish in the Salish Sea from the coastal populations. A single individual from the Washington and Oregon region genetically grouped with the Salish Sea and Puget Sound samples. The coastal Copper samples exhibit a more continuous distribution along PC2, with the most distal samples from California and Hecate Strait grouping more closely together than with most Washington and Oregon samples.

**Figure 2:**
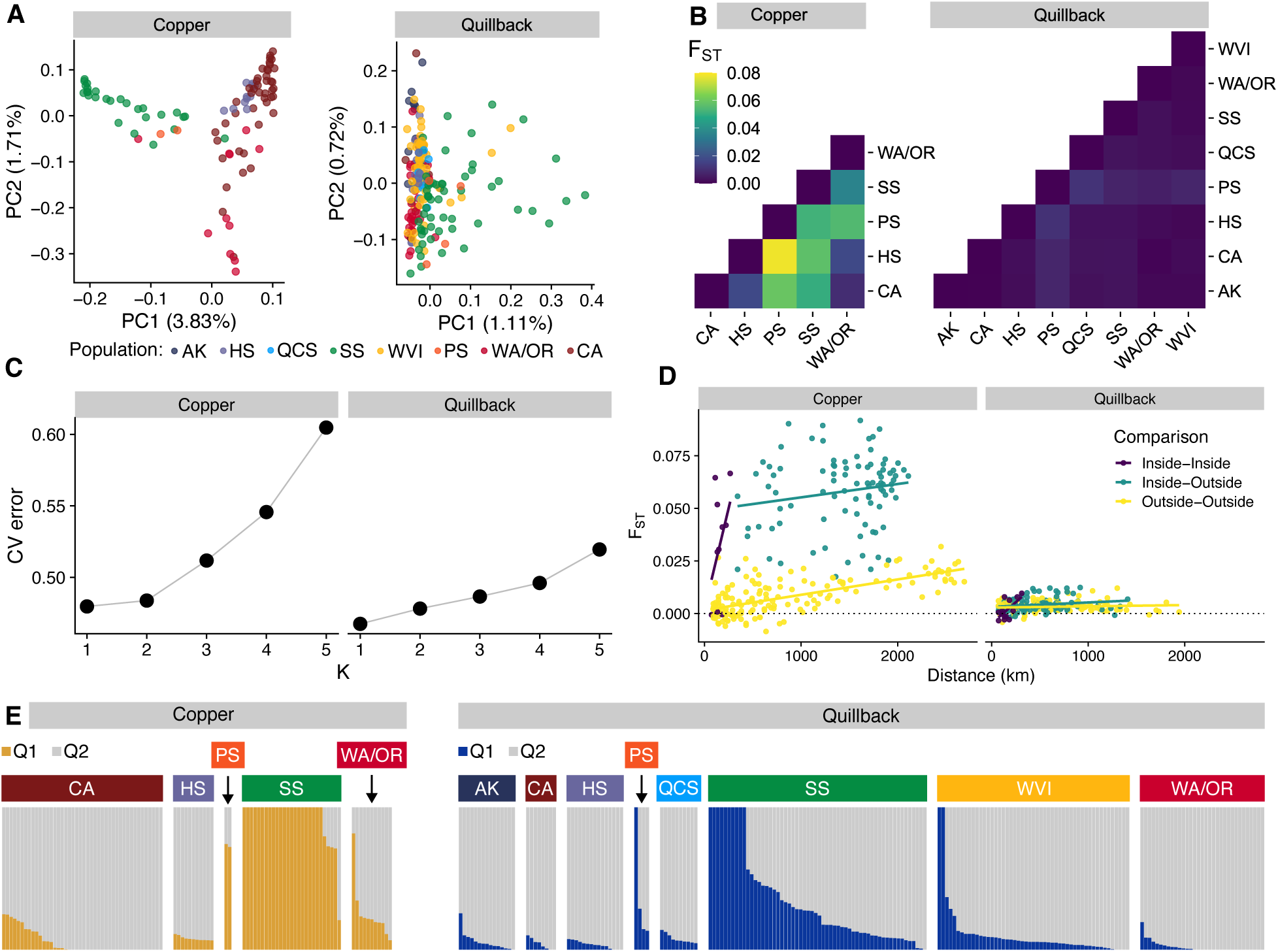
Intraspecific population structure and differentiation. **A)** Principal component analyses for both species, individual points coloured by geographic population, see legend in B. **B)** Heatmap of differentiation between geographic populations in both species. **C)** Cross-validation error for intraspecific *admixture* analyses. **D)** Genetic differentiation versus Haversine distance for coordinate populations. Inside includes Salish Sea and Puget Sound populations and outside includes all others. **E)** Intraspecific ancestry proportions as K = 2. See Figure S3 for a Quillback-only version of panels B and D.

Quillback PCAs are less informative, as most samples tend toward a single dense cluster (Fig. 2A). The first PC largely separates out a subset of Salish Sea, Puget Sound and West Vancouver Island samples, with outliers concentrating in the narrow straits at the North end of the Salish Sea (Fig. S2).

Within Copper, F_ST_ between the population bins was highest in inside (e.g. Salish Sea) vs outside (all others) comparisons (Fig. 2C). Values were highest between Puget Sound and Hecate Strait (F_ST_ = 0.078), and lowest between California and Washington/Oregon (F_ST_ = 0.0095). The pattern in Quillback was similar, but population differentiation was much lower – the highest F_ST_ value was between Queen Charlotte Sound and Puget Sound (F_ST_ = 0.011), while the lowest, highlighting the very low differentiation of coastal Quillback, was between California and Alaska, (F_ST_ = 0.0003) (Fig. 2C, Fig. S3).

Analysis of pairwise differentiation by coordinate showed a similar trend (Fig. 2D). In Copper, mean pairwise F_ST_ between coastal sampling locations was 0.008, between locations in inside waters was 0.033, and in pairwise comparisons between inside and outside waters was 0.058. In Quillback, these values were much lower: mean F_ST_ of comparisons between outside waters locations was 0.003, within inside was 0.002, and between inside and outside was 0.004.

ADMIXTURE analyses were somewhat similar between species (Fig. 2E); the highest confidence estimate of population structure (lowest cross-validation error; Fig. 2C) was in K = 1. In Copper, cross-validation error for K = 2 was similarly low and reflective of the population structure observed in the PCA. At K = 2, Copper within Puget Sound and the Salish Sea as well as a single northern Washington coast sample appear to make up one group, while other coastal populations maintain some small proportion of that ancestry. Quillback are similar but the pattern is weaker, which is consistent with the low overall levels of differentiation observed in the species.

When separating inside and outside Copper samples we see some evidence of regional structure. For example, at K = 3, Copper samples largely, although not perfectly, separate into Californian, Washington/Oregon and Hecate Strait clusters, although K = 1 has the lowest cross-validation error (Fig. S4). For inside Copper, PCA shows no patterning, but admixture suggests a separate Puget Sound cluster, again without support from CV error (Fig. S5). Outside Quillback show a subtle signal of regional clustering in PCA, but no clear patterning in admixture (Fig. S6). Finally, inside Quillback show no clustering in PCA or admixture (Fig. S7).

Both full and partial (controlling for inside vs outside comparisons) Mantel test revealed significant isolation by distance, with a moderate positive correlation between genetic and geographic distance in Copper (Full: r = 0.464, P = 0.0006, 10,000 permutations. Partial: r = 0.520, P = 0.0001, 10,000 permutations). Mantel tests were also significant when restricted to outside-outside comparisons (Full: r = 0.674, P = 0.0001, 10,000 permutations). For Quillback, all Mantel tests were positive, but non-significant (Full: r = 0.134, P = 0.175, 10,000 permutations. Partial: r = 0.152, P = 0.14, 10,000 permutations), including when limited to outside-outside comparisons (Full: r = 0.102, P = 0.248, 10,000 permutations).

### Introgression of Quillback into Copper, and of an unknown species into Quillback

A basic interspecies ADMIXTURE analysis (Fig. 1G) initially suggested bidirectional introgression between Copper and Quillback. We thus tested this using Patterson’s D to ask if “admixed” samples were genetically closer to their congener than “non-admixed” samples. As predicted, admixed Copper were closer to Quillback than non-admixed (*D* = 0.0554; p < 2.2e-16), however surprisingly non-admixed Quillback were closer to Copper than admixed Quillback (*D* = -0.0131; p < 2.23e-16). This suggests a more complex introgression scenario.

Using individual sample D-scores for Copper, we found that most individuals were slightly negative, except for Salish Sea and Puget Sound samples which were strongly positive (Fig. 3A). This pattern is consistent with Quillback introgression largely localized to the Salish Sea and Puget Sound. In contrast, Quillback individuals were generally positive, except for a few strongly negative scores from samples in Vancouver Island, the Salish Sea and Puget Sound (Fig. 3B). This suggests that there is introgression from a non-Copper source in these regions driving negative D-scores and consequently, generally positive D-scores in all other samples. We found only one sample, located in the Salish Sea, with a disproportionately high D-score indicating likely true Copper introgression into Quillback.

**Figure 3:**
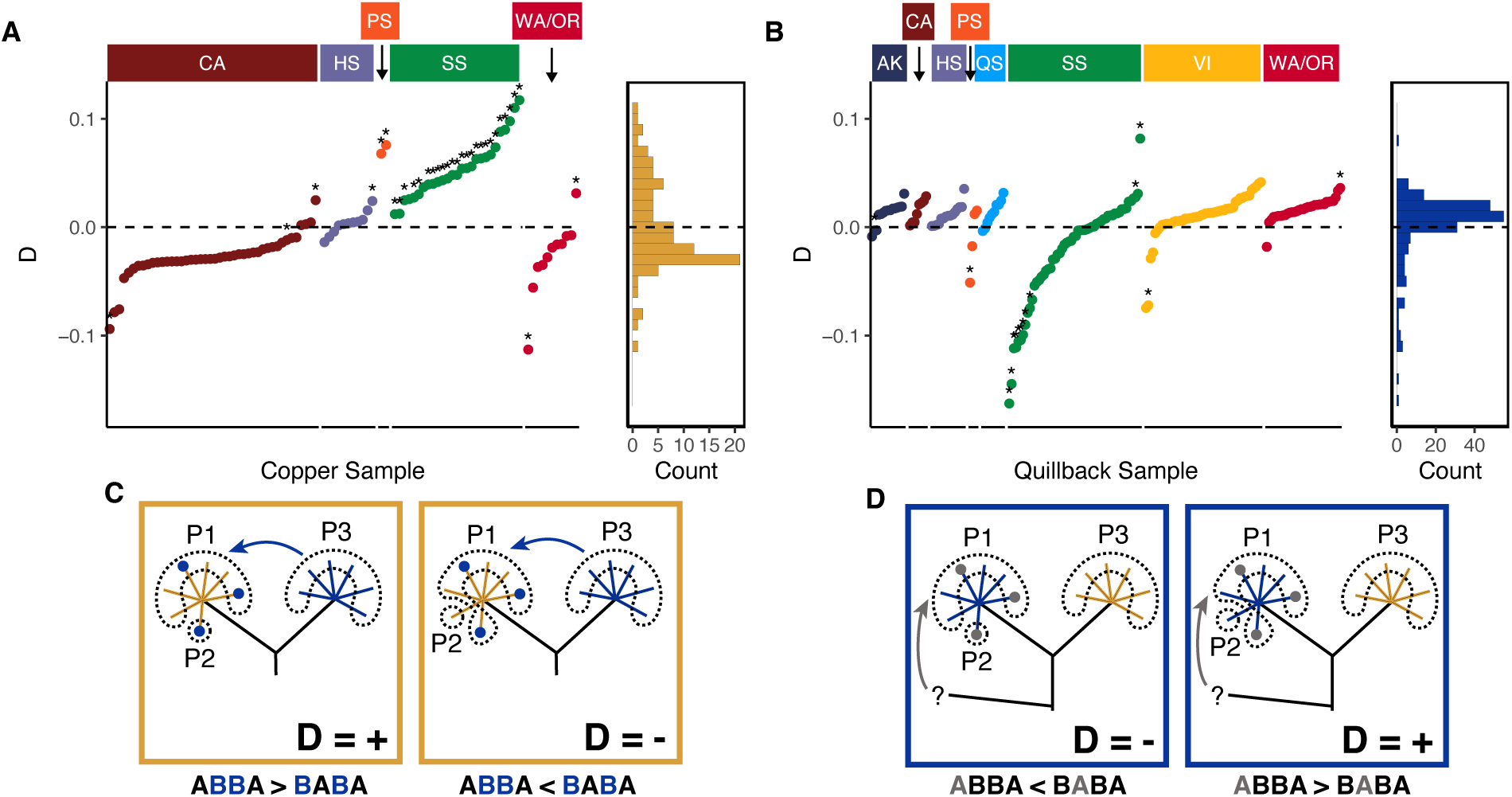
Introgression explains population structure in Rockfish. **A)** Individual D-scores for Copper testing for Quillback introgression. Individuals with > 2.5% admixture based on interspecies ADMIXTURE are highlighted. **B)** Individual D-scores for Quillback testing for Copper introgression. Individuals with > 2.5% admixture based on interspecies ADMIXTURE are highlighted. **C-D)** Conceptual diagram showing how introgression can cause positive or negative D-scores observed in A and B. Dotted lines indicate groupings for P1, P2 and P3 from the D test. Colored dots indicate admixed individuals. In Panel C, introgression shares derived alleles from the donor species (Blue) into the recipient species (Yellow) leading to positive or negative D scores depending on whether the tested P2 sample contains (+) or does not contain (-) introgression. In Panel D, introgression from a more distantly related species (Grey) contributes ancestral alleles leading to positive or negative D scores depending on whether the tested P2 sample contains (-) or does not contain (+) introgression. The donor also donates distinct derived alleles, but these are ignored by the statistic because they are not shared with the tested donor species.

### Genus-wide survey of introgression donors

Using our variant datasets including Quillback/Copper and diverse *Sebastes* species, for each sample we calculated a D-score using each species as the possible donor (P1: All other Quillback/Copper; P2: Individual Quillback/Copper sample; P3: *Sebastes* species). We interpreted the results from both datasets with the following model, described here for Quillback. If there is no admixture in any sample, all D-scores should cluster around zero without phylogenetic pattern. If a target sample is introgressed, we expect to see the highest D-score with its donor species. We also expect to see positive, but lower, D-scores for species closely related to the donor and negative D-scores for species distantly related to the donor. If the target sample is not introgressed, but other Quillback samples are, we expect to see negative D-scores with the donor species and positive D-scores with species closely related to Quillback. Since there are multiple species nearly equally related to Quillback, we expect that the positive scores should be nearly equal between multiple species. Under this framework, we measured the difference in D-score between the top three species for each target sample, effectively asking if a single species was more likely to be the donor than all other species. If there was a large gap between the first and second highest, we decided that the target sample was admixed with the top D-score as donor. If there was no gap between first and second, but a large gap between the second and third, we decided that the target sample was admixed but we could not distinguish between the top two options. We divided samples into high confidence, if the gap was greater than 0.02, and low confidence, if the gap was between 0.02 and 0.01. Additionally, we required that the potential donor to have a p-value < 0.0001 based on the D-stat jackknife bootstrap.

Under our analysis framework, we found diverse sources of introgression in both Copper and Quillback. For Copper, we corroborated our previous analysis and found that most Inside waters samples have Quillback admixture, but we also found evidence for admixture from Brown and Yelloweye (Fig. 4, Fig S9). In Outside waters, we found admixture from Aurora and Canary in California and Black in Washington/Oregon. We also found individual samples with admixture from Rougheye, Whitespeckled, Deacon, China and Boccaccio.

**Figure 4:**
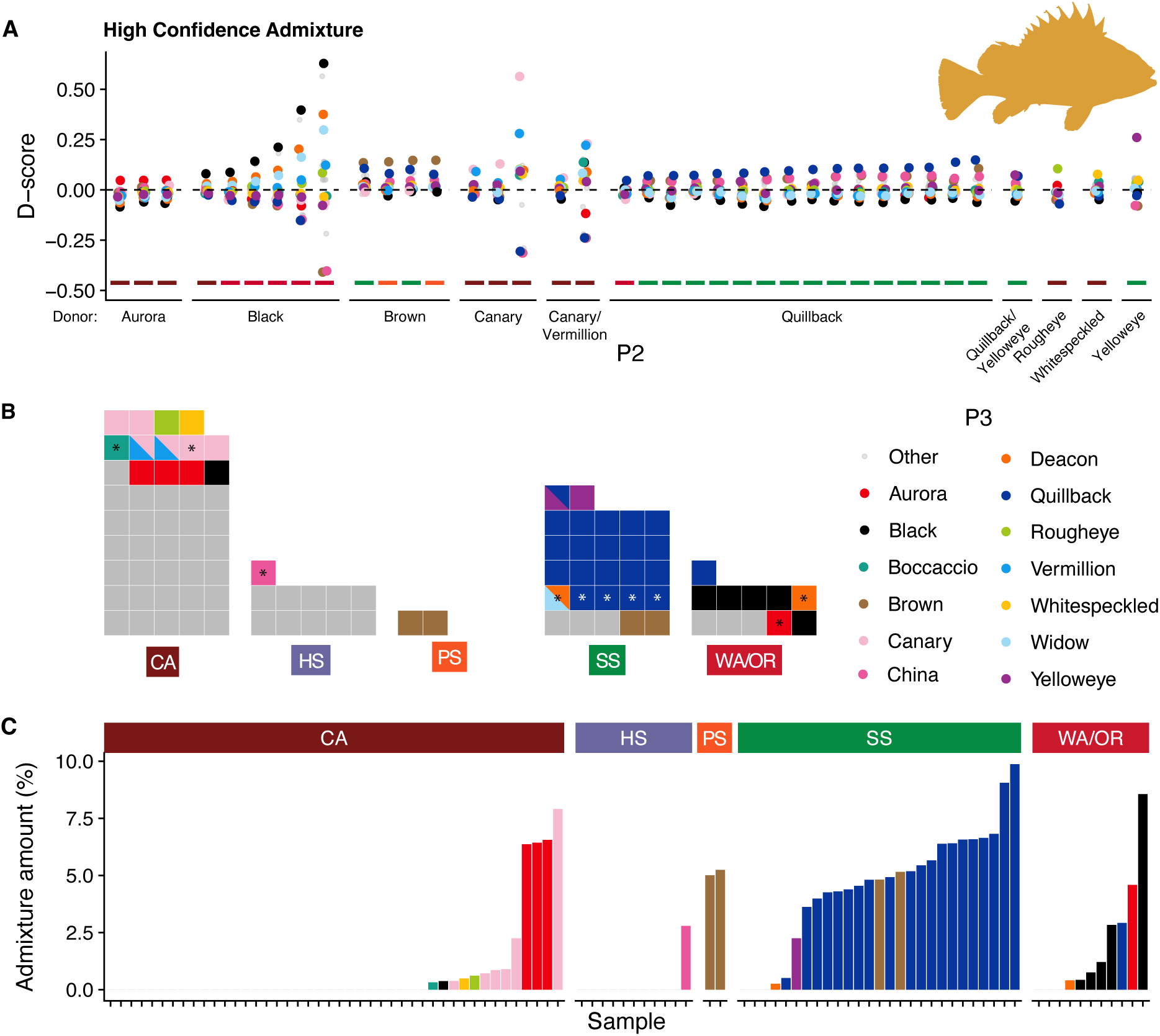
Introgression into Copper from diverse sources. A-C) D scores for Copper samples with all species as the possible donor. Each point represents the trio (All other Copper, Tested Copper, Donor species). The bar under each sample represents its geographic location (See panel C). Species not identified as being involved in admixture are coded are smaller and grey for clarity. D) Waffle plot for samples identified as admixed in each geographic location. Each square represents one sample, colored by admixture donor. Asterisks indicate low confidence of admixture. Slashes indicate multiple possible donors. Grey squares indicate samples with no admixture. E) f_4_-ratio scores for identified donor which indicates the proportion of admixed ancestry. When multiple donors were suggested, the color from the first alphabetical species was used.

For Quillback we found samples that have positive D-score for Copper as the donor, also have nearly equal positive D-score for other Quillback relatives (e.g. China, Aurora) (Fig. 5, Fig. S10). They also had a very strong distinct negative signal from Yelloweye. This is consistent with Yelloweye introgression into a subset of Quillback, and not what is expected with Copper introgression into the target sample. Supporting this, we see strong positive Yelloweye D-scores for 14 samples, primarily in the Salish Sea. There is a relationship between identified Yelloweye D-scores and Quillback PC1, suggesting that the axis of variation detected in the PCA is Yelloweye ancestry (Fig. S8). We found high-confidence signals of admixture with Aurora/Copper, Canary, Dusky/Light Dusky, Northern, Redstripe, Rougheye, Widow and Yelloweye. Additionally, we see lower-confidence admixture with Blue, Brown, China, Greenblotched/Greenspotted, and Greenstriped. These admixed samples are geographically structured. Admixed Quillback individuals with Yelloweye ancestry tend to be found in the Salish Sea and the coast of Vancouver Island, Redstripe and Widow in the coast of Vancouver Island, Dusky/Light Dusky in Alaska and Brown in Puget Sound. Overall, there is a significantly higher proportion of admixed samples in Inside waters vs Outside waters for both Copper (90% vs 33%, Welch’s Two Sample T-test, t(79.26) = 6.86, p < 0.00001). and Quillback (57% vs 13%, Welch’s Two Sample T-test, t(61.91) = 5.64, p < 0.00001).

**Figure 5:**
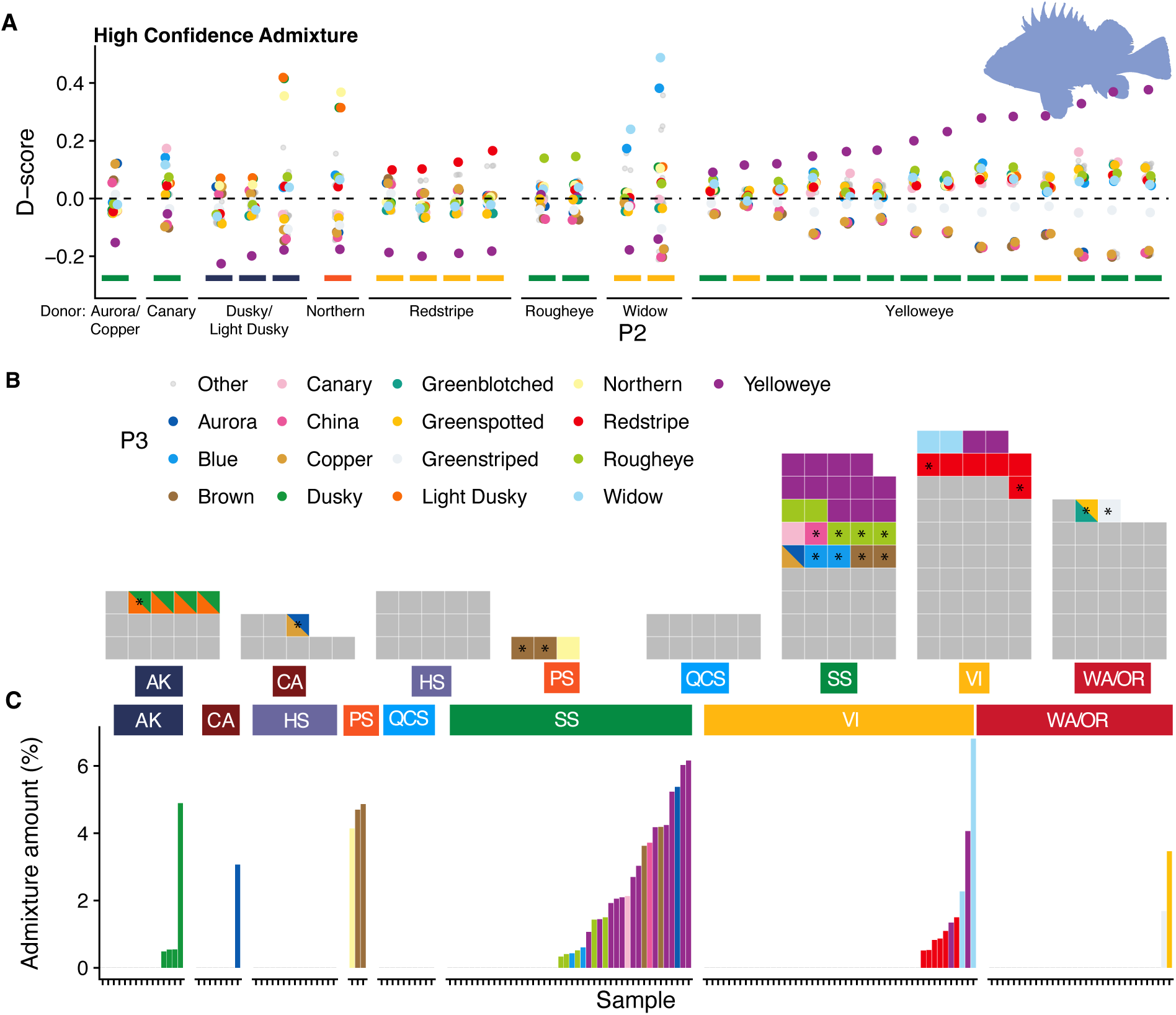
Introgression into Quillback from diverse sources. A-C) D scores for Quillback samples with all species as the possible donor. Each point represents the trio (All other Quillback, Tested Quillback, Donor species). Note that for non-admixed samples, Yelloweye had a strong negative D-score in all cases (Fig. S10). The bar under each sample represents its geographic location (See panel C). Species not identified as being involved in admixture are coded as grey. D) Waffle plot for samples identified as admixed in each geographic location. Each square represents one sample, colored by admixture donor. Asterisks indicate low confidence of admixture. Slashes indicate multiple possible donors. Grey squares indicate samples with no admixture. E) f_4_-ratio scores for identified donor which indicates the proportion of admixed ancestry. When multiple donors were suggested, the color from the first alphabetical species was used.

## Discussion

In this study, we observed significant differences in the distribution of genetic variation between Copper and Quillback rockfishes, both within the genome and between sampling locations, despite similar levels of genetic diversity. The subtle overall population structure in both species is in line with genetic and genomic analyses in other Pacific rockfish species (An et al., 2012; Buonaccorsi et al., 2004; Gao et al., 2016; Zhu et al., 2024). Higher differentiation between coast and inlet sampling locations is also consistent with previous genetic studies in this genus (Buonaccorsi et al., 2002, 2005; Dick et al., 2014), but the significant interspecific differences in range-wide population structure are here described for the first time (Fig. 6). Lastly, we showed that both Copper and Quillback have introgression from multiple sources. Interestingly, introgression from Yelloweye drove regional population structure in Quillback and introgression from Quillback contributed to structure in Copper.

**Figure 6:**
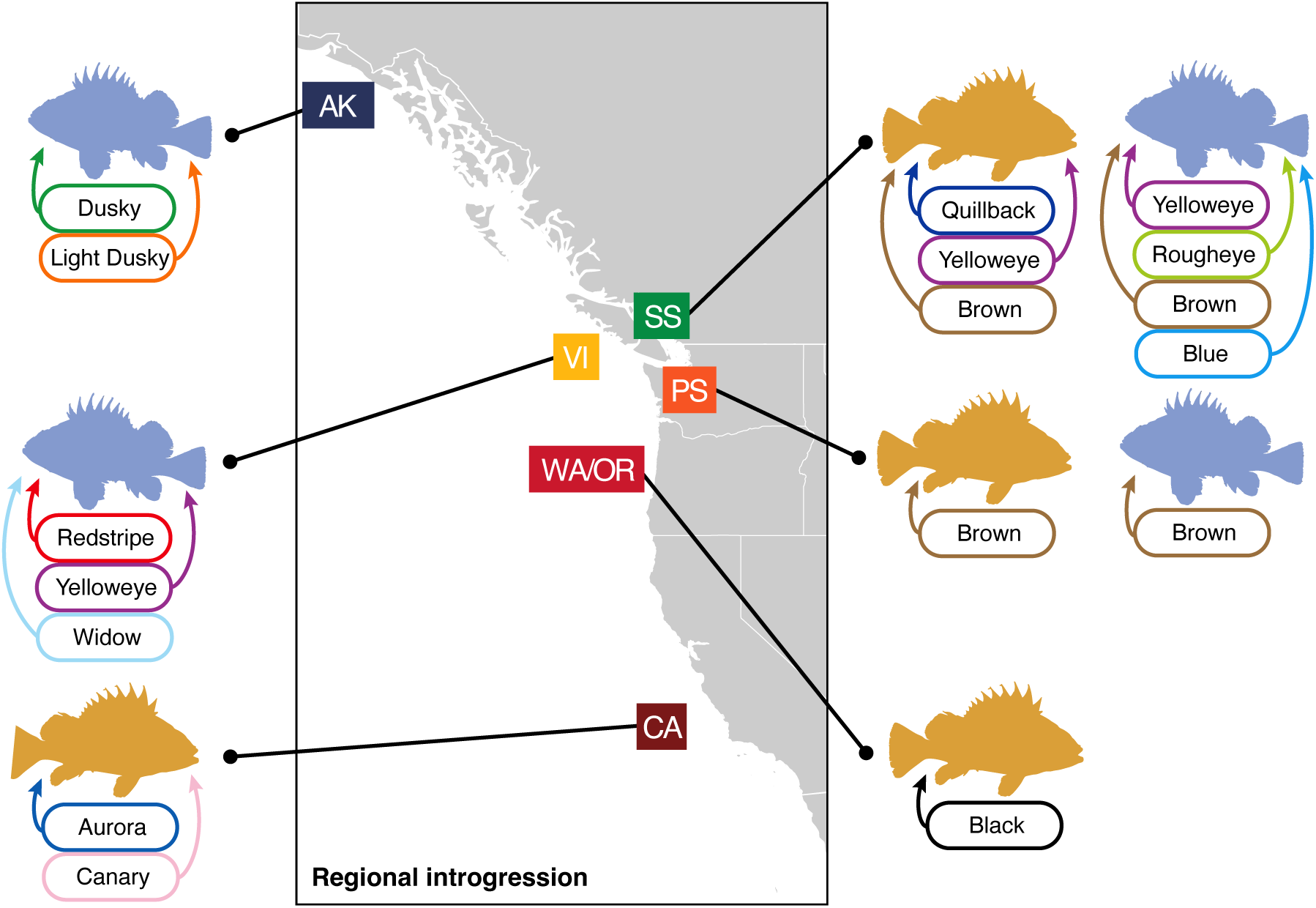
Summary of admixture in Copper and Quillback. Species highlighted are admixture donors in more than one sample in the region. Copper appears as yellow, and Quillback as blue.

### Introgression and population structure in Copper Rockfish

Hybridization is ongoing between the two target species, but introgression is largely unidirectional; genetic material primarily flows from Quillback into Copper. The low level of mixed ancestry in the sample set is consistent with long-term, low levels of hybridization, primarily restricted to the Salish Sea and Puget Sound. This largely supports previous studies using fewer markers that found introgression in the Salish Sea and Puget Sound (Seeb 1998; Schwenke et al., 2018; Wray et al., 2024). While both Schwenke et al. and Wray et al. found that Quillback admixture into Copper was focused in South Puget Sound, our analysis showed that Quillback introgression spanned the entire inside region, and there was a greater proportion of Brown introgression in Puget Sound. Unlike Quillback, we detected only a small proportion of Yelloweye introgression in the Salish Sea. Although admixture levels were higher in Inside waters, one third of Copper samples were admixed in Outside waters, suggesting that hybridization is relatively common on an evolutionary scale. The regional variation in admixture donor cannot be simply explained by range overlaps as most species have large ranges. For example, although Canary Rockfish is only an admixture donor in California, it co-occurs with Copper from Alaska to California. It’s possible that features like local population density, depth, continental slope or benthic environment lead to greater sympatry and hybridization between particular species.

Copper are significantly more differentiated throughout their range than Quillback, despite similar levels of genetic diversity. Our results support previous work suggesting that population structure of Copper rockfish in inside waters is partially driven by hybridization with Quillback (Schwenke et al., 2018; Wray et al., 2024). This means that introgression is limited to one genetic group (i.e. Salish Sea and Puget Sound), but not all members of that group have introgression. With the unique confluence of increased hybridization potential and the limitations to dispersal imposed by Salish Sea (Buonaccorsi et al., 2005), Copper rockfish exhibit a unique population structure that is amplified by directional introgression from Quillback and reinforced by the effects of genetic drift. This is not unprecedented in marine species, as local introgression has been shown to be the main cause of intraspecific differentiation in mussels (Fraïsse et al., 2016), but this finding highlights the role that introgression may play in regional genetic variation, especially in species with otherwise very low genetic differentiation.

### Introgression and population structure in Quillback Rockfish

The very low levels of genetic differentiation between Quillback populations more resemble the norm among marine fishes with long pelagic larval durations, which tend toward subtle, if any, population structure (Waples, 1998). Outside of the Salish Sea, we find some signals of subtle population structure, although all F_ST_ scores were very close to zero (0 to 0.004). The large differences in the magnitude of population structure between quillback and copper rockfish in outside waters along the Pacific coast are highly intriguing. The contrasting patterns of populations structure are unlikely the result of differences in population sizes, generation time, or PLDs as both species are relatively common and have generally similar life spans and PLDs (Love et al., 2002). Additionally, Copper and Quillback appear to occupy very similar ecological niches based on diets, growth patterns, and distributions (Murie, 1995). Juvenile Copper and Quillback also settle in similar nearshore complex habitats containing understory kelp, although they do show some subtle differences in depth preferences as adults, with Copper preferring slightly shallower water (Palsson et al., 2009). We hypothesize that cryptic ecological processes that limit the dispersal potential of Copper compared to Quillback may explain the variation in population structure that we observed but additional research is necessary to investigate this.

Initial admixture and D-score tests suggested Copper introgression in Quillback, but using a wide array of possible donors we showed that this signal was not Copper-specific and instead likely was a reverse signal from Yelloweye introgression in the Salish Sea. Yelloweye is not an expected candidate for hybridization with Quillback because they rarely co-occur, although they geographically overlap, and they are not in the same clade and diverged approximately 7 mya (Kolora et al., 2020; Love et al., 2002). Interestingly, admixture seems to be geographically structured. Yelloweye and Rougheye admixture are primarily in the Salish Sea, Widow and Redstripe admixture in the outer coast of Vancouver Island and Dusky in Alaska. Surprisingly, we see a signal of admixture with Northern Rockfish in Puget Sound, where Northern Rockfish do not occur. Since the admixture is only 4% of the genome, the admixed fish is multiple generations from the initial hybridization event and may have migrated long distances over generations. Alternatively, the Northern Rockfish signal may have originated from another closely related species and be misattributed based on the incomplete sampling of possible donors.

Overall, allele sharing appears enhanced by hybridization of opportunity or necessity in more complex and degraded estuarine habitats, but is not exclusively in these areas. The low levels of admixture we observe here could be due to temporary breakdowns of reproductive barriers. Depending on the amount of hybridization, the relative population sizes and the amount of selection on introgressed ancestry, signals of admixture can persist in a population for thousands of generations. Future work should attempt to date hybridization events and disentangle the relative genetic contributions of multiple species.

### Approaches to identifying admixture

We approached introgression using Patterson’s D because of the number of possible species involved. While other approaches attempt to disentangle multiple introgression events (e.g. TreeMix, PhyloNet), they are designed for smaller number of species and admixture events and would not function well with 51 species and more than a dozen introgression events (Hibbins and Hahn, 2022). Similarly, approaches like ADMIXTURE or Ohana work better with more reference samples and would struggle computationally with 51 possible donors (Cheng et al., 2017). Even when forcing each possible donor into its own admixture group, those groups will quickly become dominated by admixed samples and lose distinctiveness. Our approach directly highlights that significant D-scores can occur even when not using the true donor species in the test. This means it is possible that in some cases, our suggested donors are not accurate because we did not include the true donor species in our reference set. This is likely the case when the signal is split between two closely related species; the true donor may be a third unsampled species. We also recognize that the confidence values, based on the gap between top D-scores, is arbitrary. The traditional mechanism of assessing significance for D is a block jackknife, but because of the multiple introgression events, these tests are almost entirely significant and don’t reflect the true confidence in donor choice.

Our work shows how admixture from unsampled or ghost sources can influence admixture estimates. In our case, unadmixed Quillback look more like Copper because Yelloweye-admixed Quillback have extra ancestral alleles from the Yelloweye introgression, thereby comparatively increasing derived allele sharing between unadmixed Quillback and Copper. The strength of this mirror signal depends on the phylogenetic relatedness of each species involved in the test. One approach to understanding these patterns is through building admixture graphs, like in admixtools, although they work best with smaller numbers of samples or populations and admixture events (Maier & Patterson, 2024).

Future work should include as many *Sebastes* species as possible, although even with an exhaustive approach, extinct species may still leave a genetic signal and cannot be included. Our measures of introgression amount, the f_4_-ratio, assumes only one introgression event and it is not clear how all other events will influence this value. Lastly, our approach can only identify the largest introgression contributor and either ignores or is confounded by secondary introgression. For example, a balanced score between two species may be because both species have contributed equally. The numeric D-score also depends on both the admixture amount and the phylogenetic distance between donor and source, further complicating interpretation. Taken together, we think our approach is conservative and likely underestimates true admixture events. Further work could focus on likely donors identified here and use more targeted approaches to disentangle admixture patterns.

## Conclusions and management implications

Like many other inshore rockfishes, Copper and Quillback are currently managed as two separate populations in BC and WA: inside waters of the Salish Sea, and outside waters (Fisheries and Oceans Canada, 2024; NMFS, 2023). Our findings suggest that the current management regime separating inside and outside waters is likely representative of the biological reality in Copper. In terms of outside Copper populations, our work generally supports the current approach of management boundaries corresponding to state/province boundaries. For outside Quillback populations, our data indicates very little structure and near panmixia. Genetic data for Quillback were unavailable when management boundaries were constructed therefore the boundaries were constructed based on what was known in Copper. Constructing boundaries based on the closely related Copper was reasonable for Quillback at the time, but our study illustrates that the two species have substantially different patterns of population structure. Therefore, the current boundaries are not well supported based on genetic data alone but may be justified based on other factors such as demography, fisheries exploitation, or habitat availability. Our work also highlights that in species with little or no regional population structure, variation in introgression can be the major axis of variation picked up in methods like ADMIXTURE. Importantly, when the true source of introgression is not included, ADMIXTURE can misrepresent the source. Finally, our work suggests that hybridization in *Sebastes* is far broader than previously thought and can occur between species not thought to occupy similar niches or generally co-occur. Genomic-based studies can detect hybridization that occurred hundreds or thousands of years ago that leave no obvious morphological signal, thus we are getting a fuller picture of how genes have been exchanged in this rapidly speciating genus. Future studies should explore the genes that are being shared between species and their potentially adaptive role. Lastly, studies in *Sebastes* should appreciate that reproductive barriers are more porous than previously thought and a wide variety of hybrids are possible.

## Acknowledgements

We would like to thank Schon Hardy from Fisheries and Oceans Canada, and John Hyde, Krista Nichols, and Matt Craig from the National Oceanic and Atmospheric Administration for sharing tissue samples necessary for this work. We thank Katie D’Amelio and Kirby Karpan from the National Oceanic and Atmospheric Administration Alaska Fisheries Science Center for processing tissue samples and preparing libraries for whole genome sequencing. We thank Krista Nichols for helpful feedback on the manuscript. This work was supported by an NSERC Discovery grant and funding from Fisheries and Ocean Canada to GLO. This work was also supported by NIH National Institute of General Medicine award R35GM142916 to PHS, and NPRB project 2112 to PHS. Computational work was supported by the Digital Research Alliance of Canada, the Canadian Foundation for Innovation and the British Columbia Knowledge Development Fund.

## Data Availability

All sequence data generated for this project is on the SRA listed in Table S1. Additional samples used are listed in Table S2. All code used for this project is available at https://github.com/owensgl/copper_quillback

## References

Alexander, D. H., Novembre, J., & Lange, K. (2009). Fast model-based estimation of ancestry in unrelated individuals. Genome Research, 19(9), 1655–1664. 10.1101/gr.094052.109

An, H., Kim, M. J., Park, K., Cho, K., Bae, B., Kim, J., & Myeong, J. I. (2012). Genetic Diversity and Population Structure in the Heavily Exploited Korean Rockfish, *Sebastes schlegelii*, in Korea. Journal of the World Aquaculture Society, 43(1), 73–83. 10.1111/j.1749-7345.2011.00544.x

Anderson, S. C., Keppel, E. A., Edwards, A. M. 2019. A reproducible data synopsis for over 100 species of British Columbia groundfish. DFO Can. Sci. Advis. Sec. Res. Doc. 2019/041. 321 p.

Andrews, K. S., Nichols, K. M., Elz, A., Tolimieri, N., Harvey, C. J., Pacunski, R., Lowry, D., Yamanaka, K. L., & Tonnes, D. M. (2018). Cooperative research sheds light on population structure and listing status of threatened and endangered rockfish species. Conservation Genetics, 19(4), 865–878. 10.1007/s10592-018-1060-0

Andrews, S. (2010). FASTQC: A Quality Control Tool for High Throughput Sequence Data. http://www.bioinformatics.babraham.ac.uk/projects/fastqc/

Artamonova, V. S., Makhrov, A. A., Karabanov, D. P., Rolskiy, A. Y., Bakay, Y. I., & Popov, V. I. (2013). Hybridization of beaked redfish (*Sebastes mentella*) with small redfish (*Sebastes viviparus*) and diversification of redfish (Actinopterygii: Scorpaeniformes) in the Irminger Sea. Journal of natural history, 47(25-28), 1791–1801.

Baym, M., Kryazhimskiy, S., Lieberman, T. D., Chung, H., Desai, M. M., & Kishony, R. (2015). Inexpensive multiplexed library preparation for megabase-sized genomes. PloS one, 10(5), e0128036. 10.1371/journal.pone.0128036

Bernatchez, L., Wellenreuther, M., Araneda, C., Ashton, D. T., Barth, J. M. I., Beacham, T. D., Maes, G. E., Martinsohn, J. T., Miller, K. M., Naish, K. A., Ovenden, J. R., Primmer, C. R., Young Suk, H., Therkildsen, N. O., & Withler, R. E. (2017). Harnessing the Power of Genomics to Secure the Future of Seafood. Trends in Ecology & Evolution, 32(9), 665– 680. 10.1016/j.tree.2017.06.010

Broad Institute. (2019). Picard Toolkit. *Broad Institute, GitHub Repository*. https://github.com/broadinstitute/picard

Buonaccorsi, V. P., Kimbrell, C. A., Lynn, E. A., & Vetter, R. D. (2002). Population structure of Copper Rockfish (*Sebastes caurinus*) reflects postglacial colonization and contemporary patterns of larval dispersal. Canadian Journal of Fisheries and Aquatic Sciences, 59(8), 1374–1384. 10.1139/f02-101

Buonaccorsi, V. P., Westerman, M., Stannard, J., Kimbrell, C., Lynn, E., & Vetter, R. D. (2004). Molecular genetic structure suggests limited larval dispersal in Grass Rockfish, *Sebastes rastrelliger*. Marine Biology, 145(4), 779–788. 10.1007/s00227-004-1362-2

Buonaccorsi, V. P., Kimbrell, C. A., Lynn, E. A., & Vetter, R. D. (2005). Limited realized dispersal and introgressive hybridization influence genetic structure and conservation strategies for Brown Rockfish, Sebastes auriculatus. Conservation Genetics, 6(5), 697– 713. 10.1007/s10592-005-9029-1

Chen, S. (2023). Ultrafast one-pass FASTQ data preprocessing, quality control, and deduplication using fastp. iMeta, 2(2), e107. 10.1002/imt2.107

Cheng, J. Y., Mailund, T., & Nielsen, R. (2017). Fast admixture analysis and population tree estimation for SNP and NGS data. Bioinformatics, 33(14), 2148–2155.

Czech, L., & Exposito-Alonso, M. (2021). grenepipe: A flexible, scalable, and reproducible pipeline to automate variant and frequency calling from sequence read*s* (arXiv:2103.15167). arXiv. 10.48550/arXiv.2103.15167

Danecek, P., Auton, A., Abecasis, G., Albers, C. A., Banks, E., DePristo, M. A., Handsaker, R. E., Lunter, G., Marth, G. T., Sherry, S. T., McVean, G., Durbin, R., & 1000 Genomes Project Analysis Group. (2011). The variant call format and VCFtools. Bioinformatics, 27(15), 2156–2158. 10.1093/bioinformatics/btr330

Danecek, P., Bonfield, J. K., Liddle, J., Marshall, J., Ohan, V., Pollard, M. O., Whitwham, A., Keane, T., McCarthy, S. A., Davies, R. M., & Li, H. (2021). Twelve years of SAMtools and BCFtools. GigaScience, 10(2), giab008. 10.1093/gigascience/giab008

Díaz-Arce, N., Gagnaire, P.-A., Richardson, D. E., Walter III, J. F., Arnaud-Haond, S., Fromentin, J.-M., Brophy, D., Lutcavage, M., Addis, P., Alemany, F., Allman, R., Deguara, S., Fraile, I., Goñi, N., Hanke, A. R., Karakulak, F. S., Pacicco, A., Quattro, J. M., Rooker, J. R., … Rodríguez-Ezpeleta, N. (2024). Unidirectional trans-Atlantic gene flow and a mixed spawning area shape the genetic connectivity of Atlantic Bluefin Tuna. Molecular Ecology, 33(1), e17188. 10.1111/mec.17188

Dick, S., Shurin, J. B., & Taylor, E. B. (2014). Replicate divergence between and within sounds in a marine fish: The Copper Rockfish (*Sebastes caurinus*). Molecular Ecology, 23(3), 575–590. 10.1111/mec.12630

Euclide, P. T., Larson, W. A., Shi, Y., Gruenthal, K., Christensen, K. A., Seeb, J., & Seeb, L. (2023). Conserved islands of divergence associated with adaptive variation in sockeye salmon are maintained by multiple mechanisms. Molecular Ecology, n/a(n/a). 10.1111/mec.17126

Ewels, P., Magnusson, M., Lundin, S., & Käller, M. (2016). MultiQC: Summarize analysis results for multiple tools and samples in a single report. Bioinformatics, 32(19), 3047– 3048. 10.1093/bioinformatics/btw354

Fisheries and Oceans Canada (DFO). (2024). Groundfish Integrated Fisheries Management Plan 2024. https://waves-vagues.dfo-mpo.gc.ca/library-bibliotheque/41255732.pdf

Fraïsse, C., Belkhir, K., Welch, J. J., & Bierne, N. (2016). Local interspecies introgression is the main cause of extreme levels of intraspecific differentiation in mussels. Molecular Ecology, 25(1), 269–286. 10.1111/mec.13299

Gao, T., Han, Z., Zhang, X., Luo, J., Yanagimoto, T., & Zhang, H. (2016). Population genetic differentiation of the Black Rockfish *Sebastes schlegelii* revealed by microsatellites. Biochemical Systematics and Ecology, 68, 170–177. 10.1016/j.bse.2016.07.013

Garrison, E., & Marth, G. (2012). *Haplotype-based variant detection from short-read sequencing* (arXiv:1207.3907). arXiv. 10.48550/arXiv.1207.3907

Gilbert-Horvath, E. A., Larson, R. J., & Garza, J. C. (2006). Temporal recruitment patterns and gene flow in Kelp Rockfish (*Sebastes atrovirens*). Molecular Ecology, 15(12), 3801– 3815. 10.1111/j.1365-294X.2006.03033.x

Hannah, R. W., & Rankin, P. S. (2011). Site Fidelity and Movement of Eight Species of Pacific Rockfish at a High-Relief Rocky Reef on the Oregon Coast. North American Journal of Fisheries Management, 31(3), 483–494. 10.1080/02755947.2011.591239

He, Y., Chang, Y., Bao, L., Yu, M., Li, R., Niu, J., … & Qi, J. (2019). A chromosome-level genome of black rockfish, Sebastes schlegelii, provides insights into the evolution of live birth. Molecular ecology resources, 19(5), 1309–1321. 10.1111/1755-0998.13034

Helmstetter, A. J., Papadopulos, A. S. T., Igea, J., Van Dooren, T. J. M., Leroi, A. M., & Savolainen, V. (2016). Viviparity stimulates diversification in an order of fish. Nature Communications, 7(1), 11271. 10.1038/ncomms11271

Hibbins, M. S., & Hahn, M. W. (2022). Phylogenomic approaches to detecting and characterizing introgression. Genetics, 220(2), iyab173. 10.1093/genetics/iyab173

Hyde, J. R., & Vetter, R. D. (2007). The origin, evolution, and diversification of rockfishes of the genus *Sebastes* (Cuvier). Molecular Phylogenetics and Evolution, 44(2), 790–811. 10.1016/j.ympev.2006.12.026

Jay, P., Whibley, A., Frézal, L., Rodríguez de Cara, M. Á., Nowell, R. W., Mallet, J., Dasmahapatra, K. K., & Joron, M. (2018). Supergene Evolution Triggered by the Introgression of a Chromosomal Inversion. Current Biology, 28(11), 1839–1845.e3. 10.1016/j.cub.2018.04.072

Jun, G., Wing, M. K., Abecasis, G. R., & Kang, H. M. (2015). An efficient and scalable analysis framework for variant extraction and refinement from population scale DNA sequence data. Genome Research, gr.176552.114. 10.1101/gr.176552.114

Kolora, S. R. R., Owens, G. L., Vazquez, J. M., Stubbs, A., Chatla, K., Jainese, C., … & Sudmant, P. H. (2021). Origins and evolution of extreme life span in Pacific Ocean rockfishes. Science, 374(6569), 842–847. 10.1126/science.abg5332

Korunes, K. L., & Samuk, K. (2021). *pixy*: Unbiased estimation of nucleotide diversity and divergence in the presence of missing data. Molecular Ecology Resources, 21(4), 1359– 1368. 10.1111/1755-0998.13326

Li, H., & Ralph, P. (2019). Local PCA Shows How the Effect of Population Structure Differs Along the Genome. Genetics, 211(1), 289–304. 10.1534/genetics.118.301747

Longo, G. C., Lam, L., Basnett, B., Samhouri, J., Hamilton, S., Andrews, K., Williams, G., Goetz, G., McClure, M., & Nichols, K. M. (2020). Strong population differentiation in lingcod (*Ophiodon elongatus*) is driven by a small portion of the genome. Evolutionary Applications, 13(10), 2536–2554. 10.1111/eva.13037

Love, M. S., Yoklavich, M., & Thorsteinson, L. K. (2002). *The Rockfishes of the Northeast Pacific*. Univeristy of California Press.

Maier, R., & Patterson, N. (2024). admixtools: Inferring demographic history from genetic data. R package version 2.0.4, https://github.com/uqrmaie1/admixtools.

Malinsky, M., Matschiner, M., & Svardal, H. (2021). Dsuite—Fast D-statistics and related admixture evidence from VCF files. Molecular Ecology Resources, 21(2), 584–595. 10.1111/1755-0998.13265

Matthews, K. R. (1990). An experimental study of the habitat preferences and movement patterns of Copper, Quillback, and Brown rockfishes (*Sebastes* spp.). Environmental Biology of Fishes, 29(3), 161–178. 10.1007/BF00002217

Miller, J. A., & Shanks, A. L. (2004). Evidence for limited larval dispersal in black rockfish (*Sebastes melanops*): Implications for population structure and marine-reserve design. Canadian Journal of Fisheries and Aquatic Sciences, 61(9), 1723–1735. 10.1139/f04-111

Minh, B. Q., Nguyen, M. A. T., & von Haeseler, A. (2013). Ultrafast Approximation for Phylogenetic Bootstrap. Molecular Biology and Evolution, 30(5), 1188–1195. 10.1093/molbev/mst024

Minh, B. Q., Schmidt, H. A., Chernomor, O., Schrempf, D., Woodhams, M. D., von Haeseler, A., & Lanfear, R. (2020). IQ-TREE 2: New Models and Efficient Methods for Phylogenetic Inference in the Genomic Era. Molecular Biology and Evolution, 37(5), 1530–1534. 10.1093/molbev/msaa015

Murie, D. J. (1995). Comparative feeding ecology of two sympatric rockfish congeners, Sebastes caurinus (copper rockfish) and S. maliger (quillback rockfish). Marine Biology, 124(3), 341–353.

Muto, N., Kai, Y., Noda, T., & Nakabo, T. (2013). Extensive hybridization and associated geographic trends between two rockfishes *Sebastes vulpes* and S. *zonatus* (Teleostei: Scorpaeniformes: Sebastidae). Journal of Evolutionary Biology, 26(8), 1750–1762. 10.1111/jeb.12175

Narum, S. R., Horn, R., Willis, S., Koch, I., & Hess, J. (2023). Genetic variation associated with adult migration timing in lineages of Steelhead and Chinook Salmon in the Columbia River. Evolutionary Applications, 17(2), e13626. 10.1111/eva.13626

Natasha, J., Stockwell, B. L., Marie, A. D., Hampton, J., Smith, N., Nicol, S., & Rico, C. (2022). No Population Structure of Bigeye Tunas (*Thunnus obesus*) in the Western and Central Pacific Ocean Indicated by Single Nucleotide Polymorphisms. Frontiers in Marine Science, 9. 10.3389/fmars.2022.799684

Nielsen, E. E., Hemmer-Hansen, J., Larsen, P. F., & Bekkevold, D. (2009). Population genomics of marine fishes: Identifying adaptive variation in space and time. Molecular Ecology, 18(15), 3128–3150. 10.1111/j.1365-294X.2009.04272.x

NMFS. 2023. Magnuson-stevens act provisions; fisheries off west coast states; pacific coast groundfish fishery management plan; amendment 31. 88 Fed. Reg. 57,400 (Aug. 23, 2023). Available from https://www.govinfo.gov/content/pkg/FR-2023-08-23/pdf/2023-18089.pdf.

Okonechnikov, K., Conesa, A., & García-Alcalde, F. (2016). Qualimap 2: Advanced multi-sample quality control for high-throughput sequencing data. Bioinformatics, 32(2), 292–294. 10.1093/bioinformatics/btv566

Oksanen, J., Blanchet, F. G., Kindt, R., Legendre, P., Minchin, P. R., O’hara, R. B., … & Oksanen, M. J. (2013). Package ‘vegan’. Community ecology package, version, 2(9), 1–295.

Palsson, W. A., Tsou, T. S., Bargmann, G. G., Buckley, R. M., West, J. E., Mills, M. L., … & Pacunski, R. E. (2009). The biology and assessment of rockfishes in Puget Sound. Washington Department of Fish and Wildlife Report FPT-09-04.

Purcell, S., Neale, B., Todd-Brown, K., Thomas, L., Ferreira, M. A. R., Bender, D., Maller, J., Sklar, P., de Bakker, P. I. W., Daly, M. J., & Sham, P. C. (2007). PLINK: A Tool Set for Whole-Genome Association and Population-Based Linkage Analyses. The American Journal of Human Genetics, 81(3), 559–575. 10.1086/519795

R Core Team. *R: A language and environment for statistical computing*. R Foundation for Statistical Computing, Vienna, Austria. https://www.R-project.org/

Samuk, K., Owens, G. L., Delmore, K. E., Miller, S. E., Rennison, D. J., & Schluter, D. (2017). Gene flow and selection interact to promote adaptive divergence in regions of low recombination. Molecular Ecology, 26(17), 4378–4390. 10.1111/mec.14226

Schwenke, P. L., Park, L. K., & Hauser, L. (2018). Introgression among three rockfish species (*Sebastes* spp.) in the Salish Sea, northeast Pacific Ocean. PLOS ONE, 13(3), e0194068. 10.1371/journal.pone.0194068

Seeb, L. (1998). Gene flow and introgression within and among three species of rockfishes, *Sebastes auriculatus*, S. caurinus, and S. maliger. Journal of Heredity, 89(5), 393–403. 10.1093/jhered/89.5.393

Siegle, M. R., Taylor, E. B., Miller, K. M., Withler, R. E., & Yamanaka, K. L. (2013). Subtle Population Genetic Structure in Yelloweye Rockfish (*Sebastes ruberrimus*) Is Consistent with a Major Oceanographic Division in British Columbia, Canada. PLOS ONE, 8(8), e71083. 10.1371/journal.pone.0071083

Sokal, R. R. (1979). Testing statistical significance of geographic variation patterns. Systematic zoology, 28(2), 227–232. 10.2307/2412528

Tange, O. (2023, May 22). GNU Parallel 20230522 (’Charles’). Zenodo, 42–47.

Tarpey, C. M., Seeb, J. E., McKinney, G. J., Templin, W. D., Bugaev, A., Sato, S., & Seeb, L. W. (2018). Single-nucleotide polymorphism data describe contemporary population structure and diversity in allochronic lineages of Pink salmon (*Oncorhynchus gorbuscha*). Canadian Journal of Fisheries and Aquatic Sciences, 75(6), 987–997. 10.1139/cjfas-2017-0023

Therkildsen, N. O., & Palumbi, S. R. (2017). Practical low-coverage genomewide sequencing of hundreds of individually barcoded samples for population and evolutionary genomics in nonmodel species. Molecular ecology resources, 17(2), 194–208. 10.1111/1755-0998.12593

Tolimieri, N., Andrews, K., Williams, G., Katz, S., & Levin, P. S. (2009). Home range size and patterns of space use by lingcod, Copper rockfish and Quillback rockfish in relation to diel and tidal cycles. Marine Ecology Progress Series, 380, 229–243. 10.3354/meps07930

Tovar Verba, J., Stow, A., Bein, B., Pennino, M. G., Lopes, P. F. M., Ferreira, B. P., Mortier, M., Maia Queiroz Lima, S., & Pereira, R. J. (2022). Low population genetic structure is consistent with high habitat connectivity in a commercially important fish species (*Lutjanus jocu*). Marine Biology, 170(1), 5. 10.1007/s00227-022-04149-1

Tricou, T., Tannier, E., & de Vienne, D. M. (2022). Ghost lineages highly influence the interpretation of introgression tests. Systematic Biology, 71(5), 1147–1158. 10.1093/sysbio/syac011

Vasimuddin, Md., Misra, S., Li, H., & Aluru, S. (2019). Efficient Architecture-Aware Acceleration of BWA-MEM for Multicore Systems. 2019 IEEE International Parallel and Distributed Processing Symposium (IPDPS), 314–324. 10.1109/IPDPS.2019.00041

Wall, L., Christiansen, T., & Orwant, J. (2020). Perl v5.32.1. https://www.perl.org

Waples, R. (1998). Separating the wheat from the chaff: Patterns of genetic differentiation in high gene flow species. Journal of Heredity, 89(5), 438–450. 10.1093/jhered/89.5.438

Wray, A., Petrou, E., Nichols, K. M., Pacunski, R., LeClair, L., Andrews, K. S., Kardos, M., & Hauser, L. (2024). Contrasting effect of hybridization on genetic differentiation in three rockfish species with similar life history. Evolutionary Applications, 17(7), e13749. 10.1111/eva.13749

Xuereb, A., Rougemont, Q., Dallaire, X., Moore, J.-S., Normandeau, E., Bougas, B., Perreault-Payette, A., Koop, B. F., Withler, R., Beacham, T., & Bernatchez, L. (2022). Re-evaluating Coho salmon (*Oncorhynchus kisutch*) conservation units in Canada using genomic data. Evolutionary Applications, 15(11), 1925–1944. 10.1111/eva.13489

Zabel, R. W., Levin, P. S., Tolimieri, N., & Mantua, N. J. (2011). Interactions between climate and population density in the episodic recruitment of bocaccio, *Sebastes paucispinis*, a Pacific rockfish. Fisheries Oceanography, 20(4), 294–304. 10.1111/j.1365-2419.2011.00584.x

Zheng, X., Levine, D., Shen, J., Gogarten, S. M., Laurie, C., & Weir, B. S. (2012). A high-performance computing toolset for relatedness and principal component analysis of SNP data. Bioinformatics, 28(24), 3326–3328. 10.1093/bioinformatics/bts606

Zhu, B., Gao, T., He, Y., Qu, Y., & Zhang, X. (2024). Population Genomics of Commercial Fish *Sebastes schlegelii* of the Bohai and Yellow Seas (China) Using a Large SNP Panel from GBS. Genes, 15(5), Article 5. 10.3390/genes15050534

