## Supplementary Figures for "Phylogenetically diverse introgression drives subtle population structure in Pacific rockfishes"

Nathan T. B. Sykes[1]

R. Nicolas Lou[2]

Matthew R. Siegle[3]

Peter H. Sudmant[2]

Wesley A. Larson[4]

Gregory L. Owens[1*]

1. Department of Biology, University of Victoria, BC, Canada

2. Department of Integrative Biology, University of California, Berkeley, CA, USA

3. Fisheries and Oceans Canada, Nanaimo, BC, Canada

4. National Oceanic and Atmospheric Administration, Juneau, AK, USA

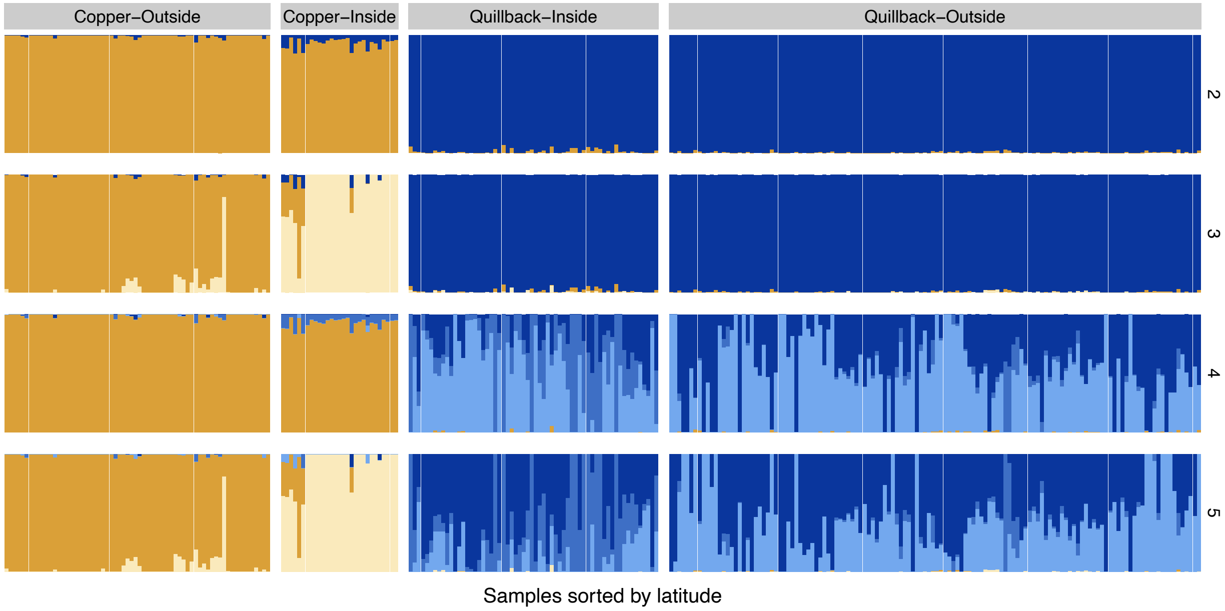


**Figure S1: Interspecies ADMIXTURE for K= 2 to 5**.

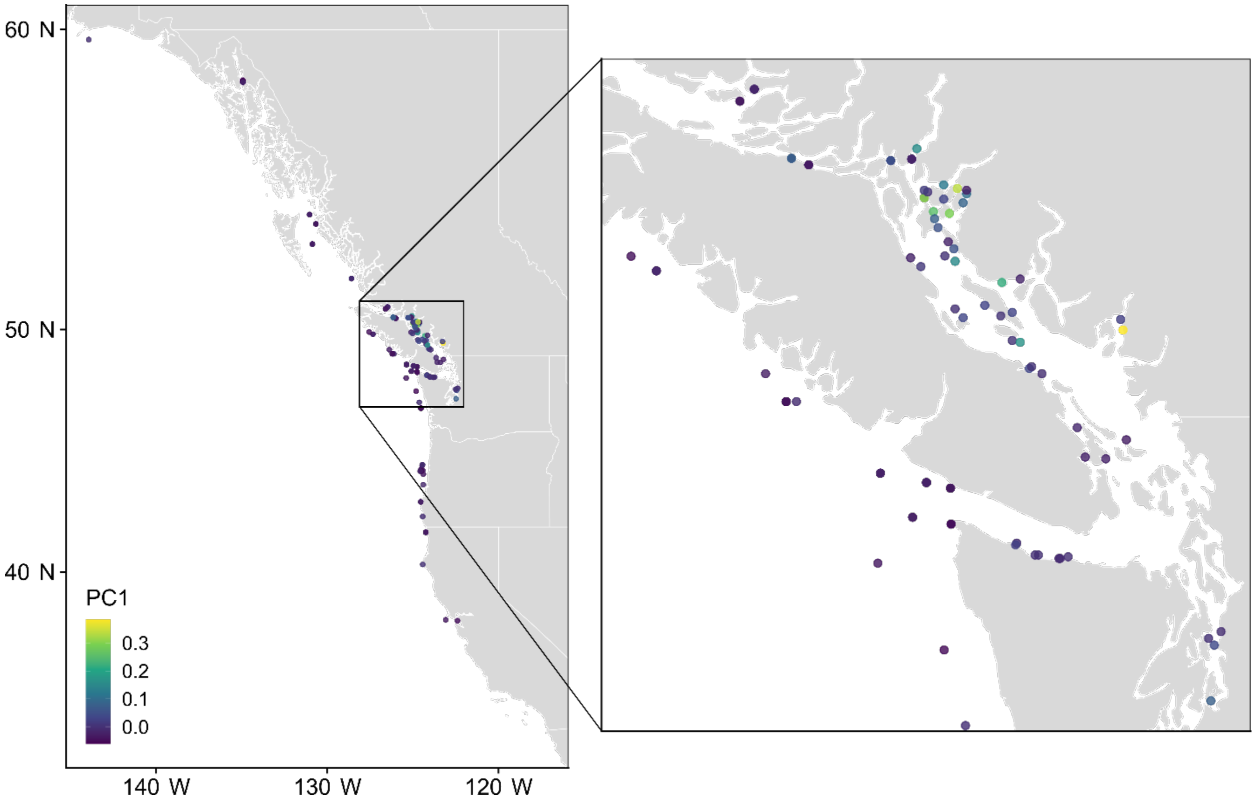


**Figure S2: Sampling map of Quillback rockfish, coloured by PC1.** PC1 outliers appear to be concentrated at the North end of the Strait of Georgia.

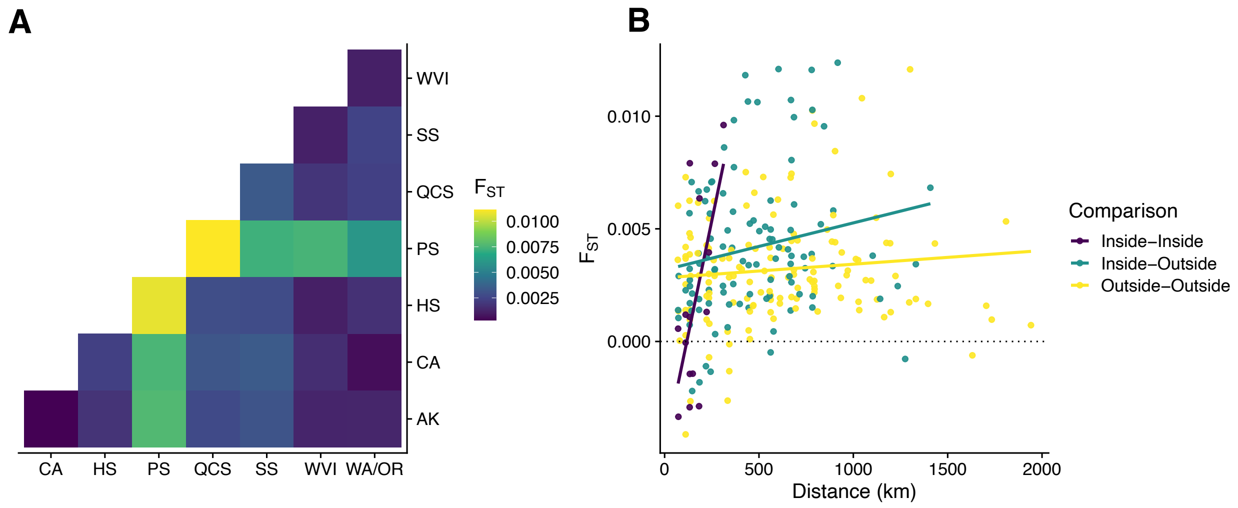


**Figure S3: $F_{ST}$ in Quillback.** A) A heatmap of $F_{ST}$ between regions. B) $F_{ST}$ between geographic coordinates, with linear trendline added for visualization.

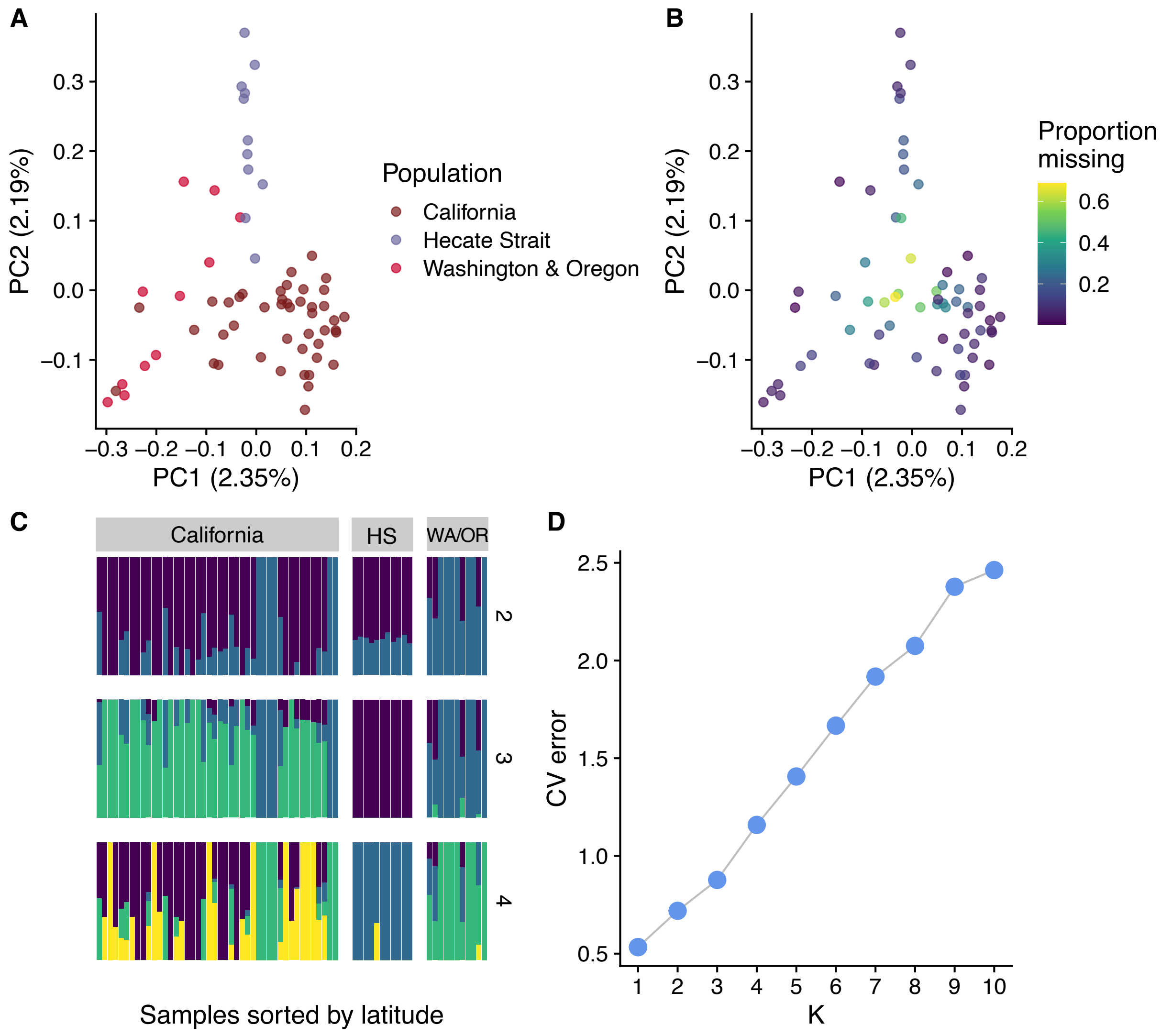


**Figure S4: PCA and ancestry analysis for Copper Rockfish outside of the Salish Sea and Puget Sound.** A) The first two principal components colored by sample region. B) The first two principal components colored by proportion of missing data. C) Samples in the ancestry plot are ordered by increasing latitude. D) Cross-validation error from admixture.

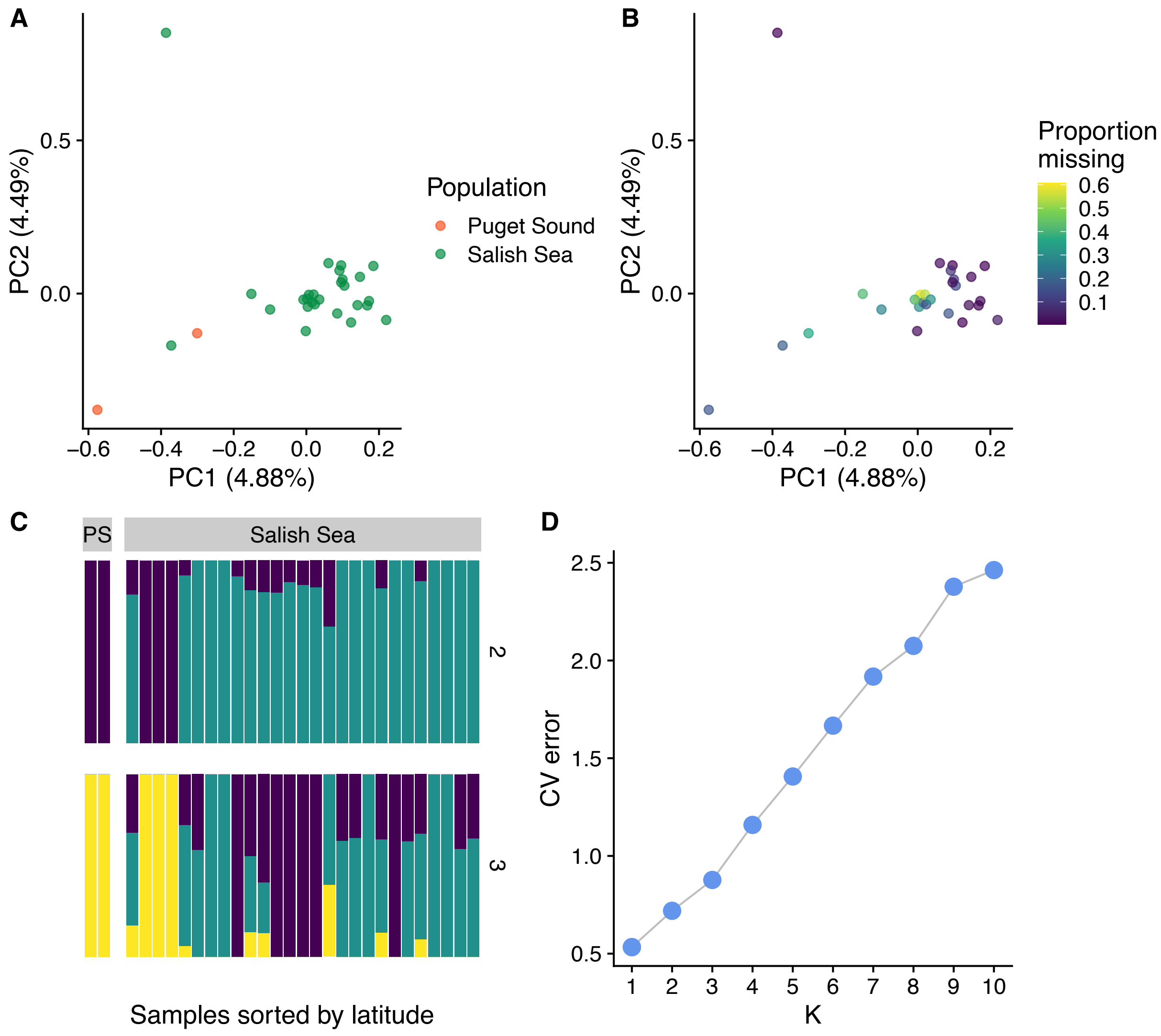


**Figure S5: PCA and ancestry analysis for Copper Rockfish inside of the Salish Sea and Puget Sound.** A) The first two principal components colored by sample region. B) The first two principal components colored by proportion of missing data. C) Samples in the ancestry plot are ordered by increasing latitude. D) Cross-validation error from admixture.

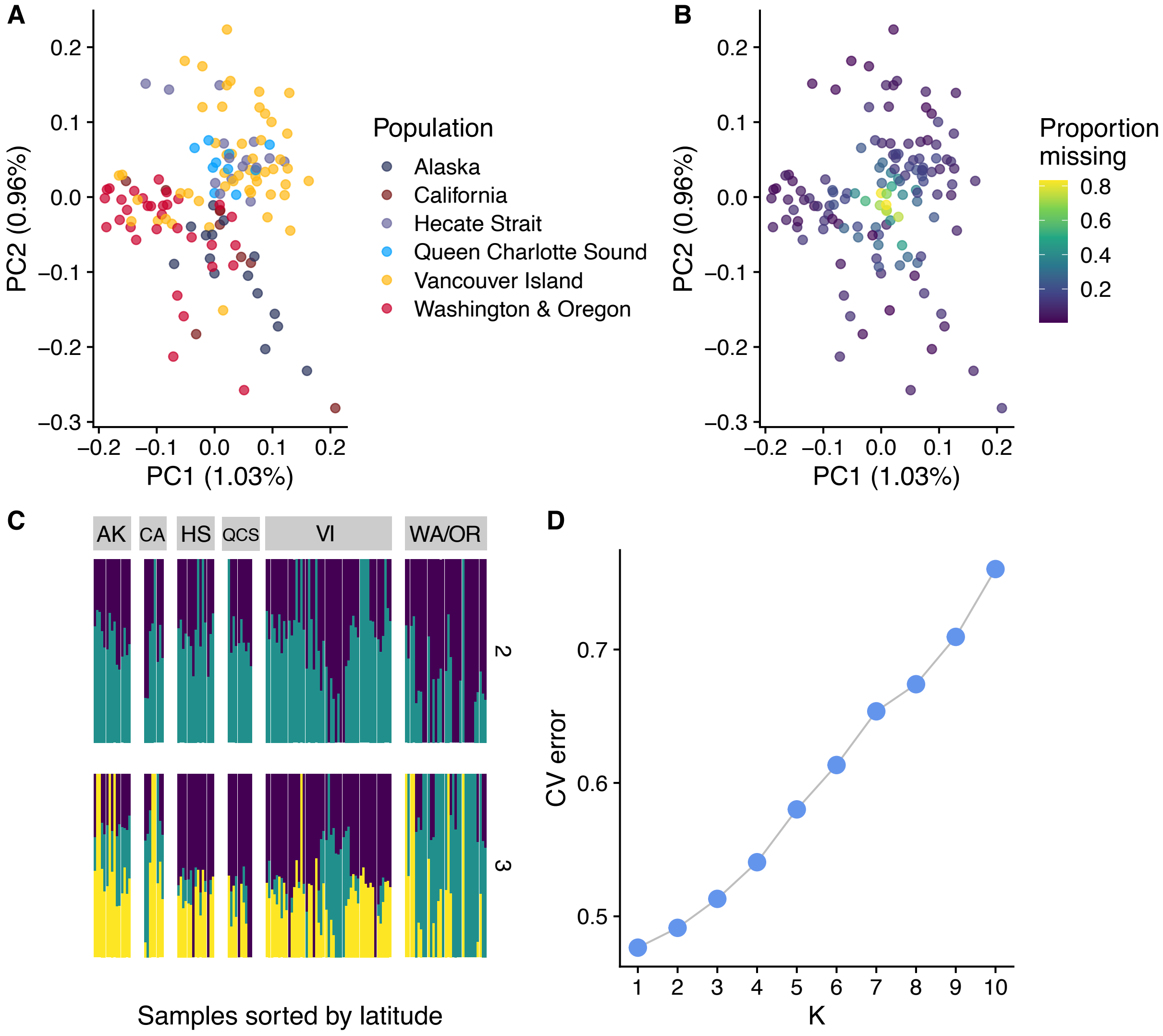


**Figure S6: PCA and ancestry analysis for Quillback Rockfish outside of the Salish Sea and Puget Sound.** A) The first two principal components colored by sample region. B) The first two principal components colored by proportion of missing data. C) Samples in the ancestry plot are ordered by increasing latitude. D) Cross-validation error from admixture.

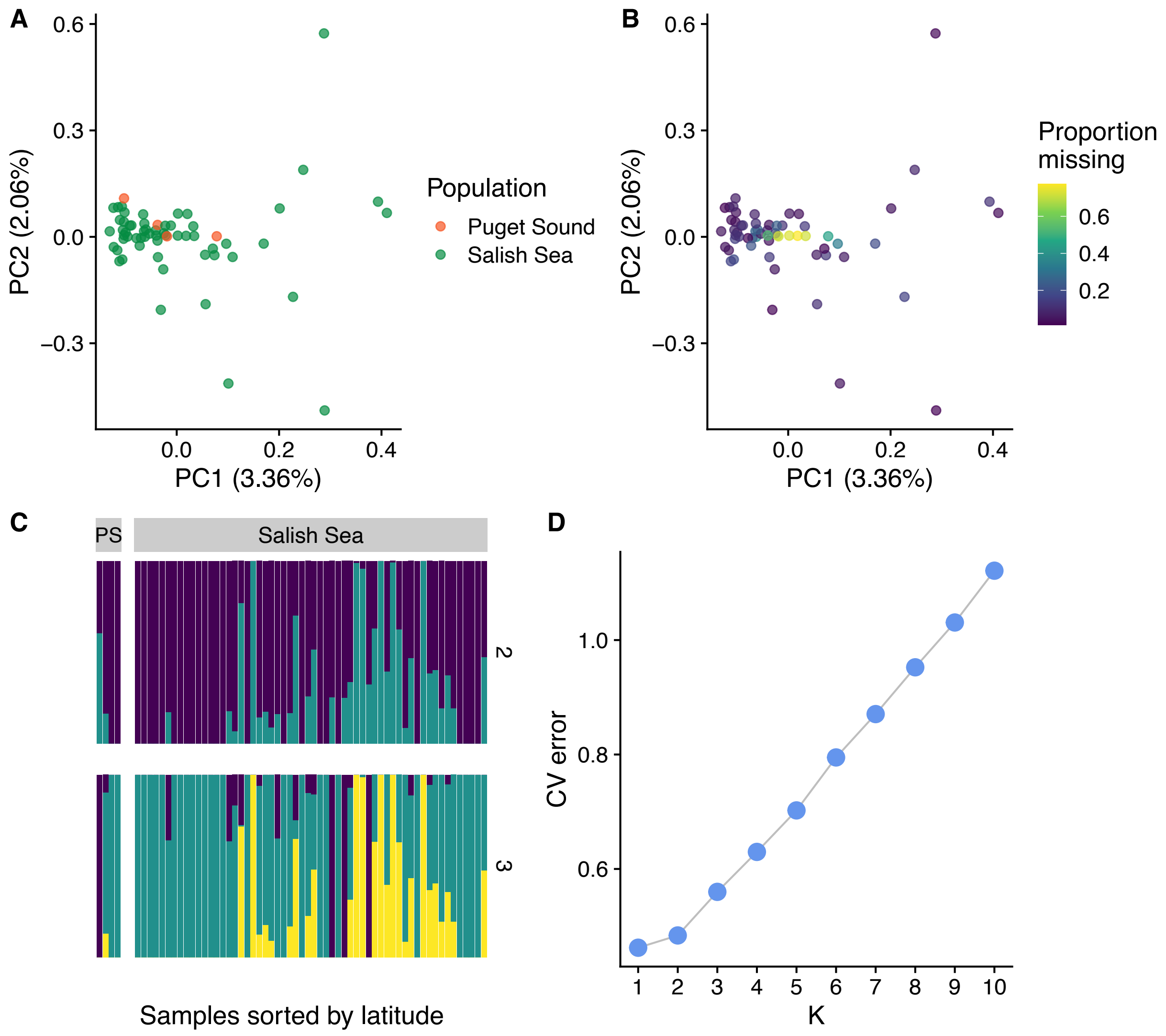


**Figure S7: PCA and ancestry analysis for Quillback Rockfish inside of the Salish Sea and Puget Sound.** A) The first two principal components colored by sample region. B) The first two principal components colored by proportion of missing data. C) Samples in the ancestry plot are ordered by increasing latitude. D) Cross-validation error from admixture.

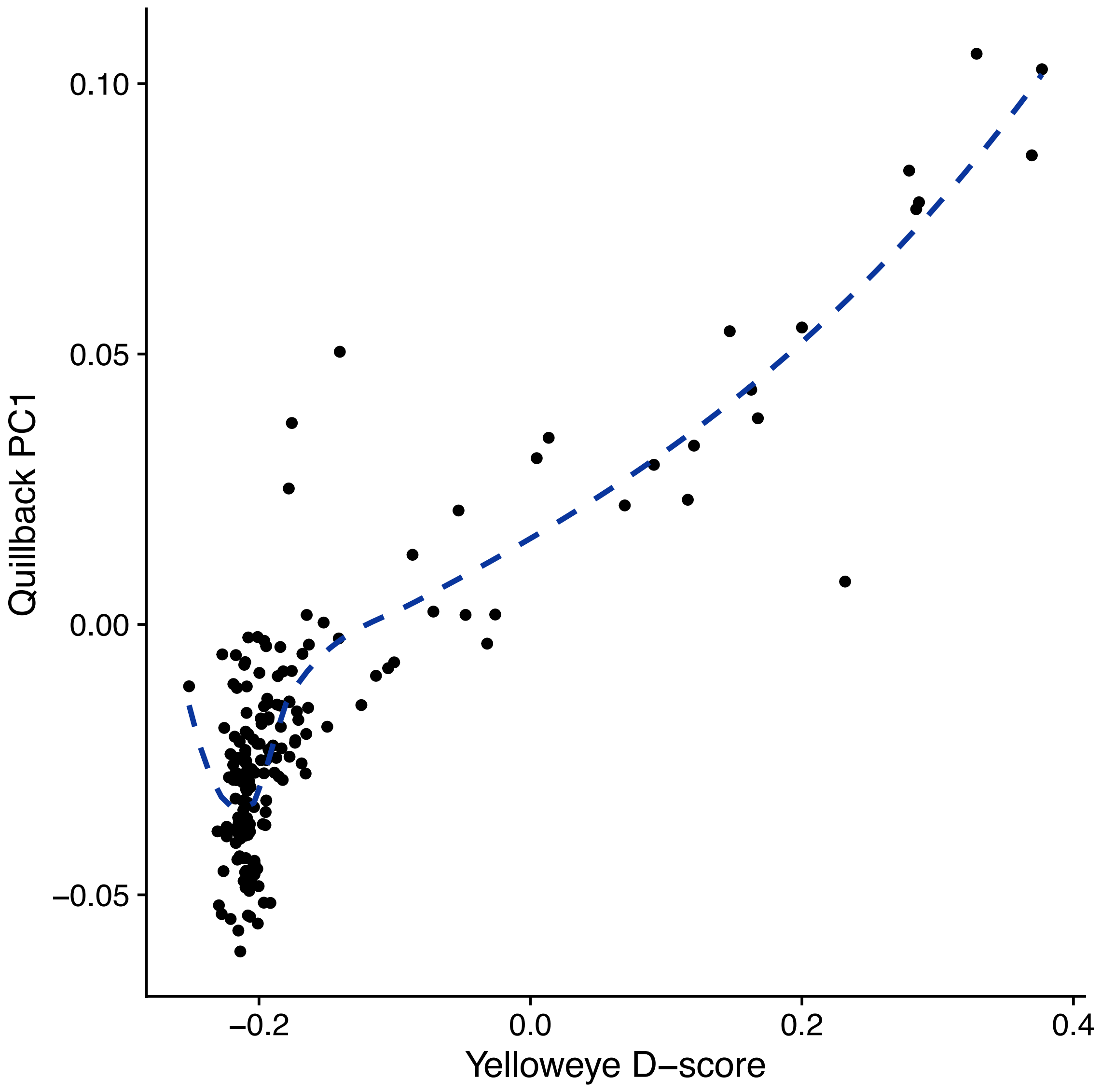


**Figure S8: Yelloweye Rockfish introgression explains Quillback PC1**

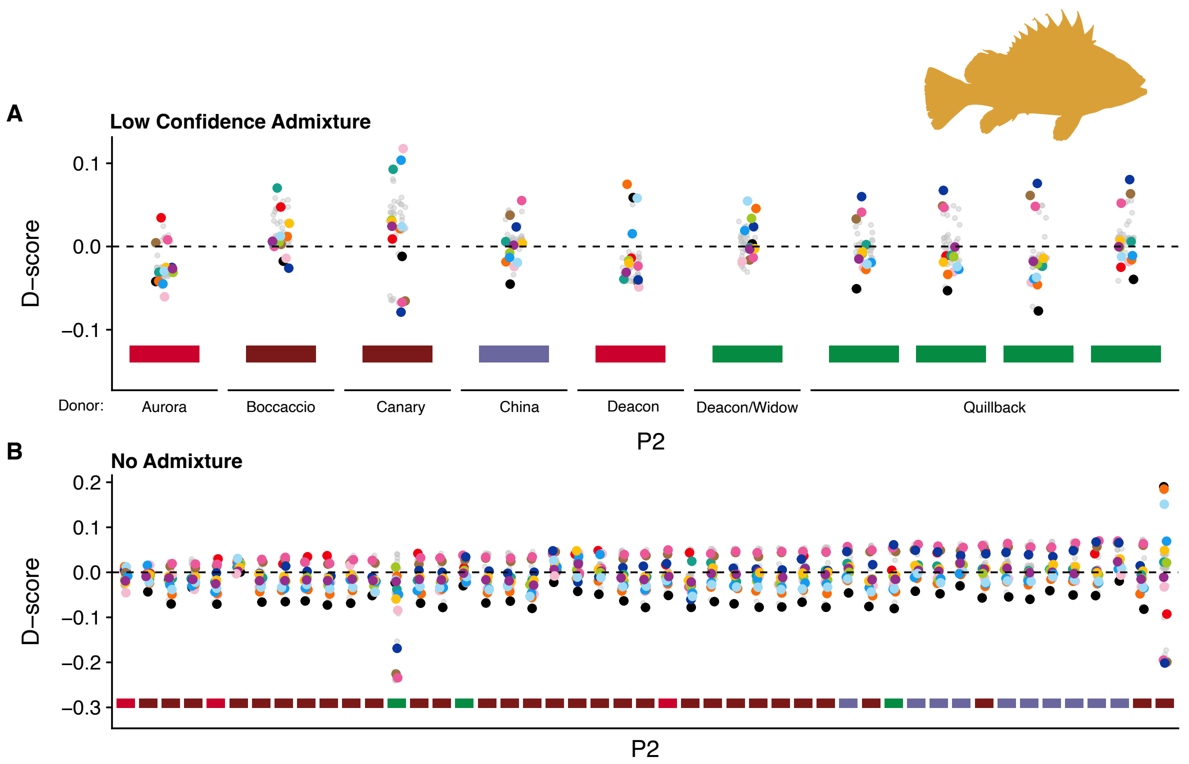


**Figure S9: Introgression into Copper from diverse sources.** D scores for Copper samples with all species as the possible donor. Each point represents the trio (All other Quillback, Tested Quillback, Donor species). The bar under each sample represents its geographic location (See Figure 2). Species not identified as being involved in admixture are coded are smaller and grey for clarity. A) Low confidence admixture: where the gap between the 1st and 2nd ranked D-score is between 0.01 and 0.02. B) No admixture: where the gap between the 1st and 2nd ranked D-score is less than 0.01, or the top admixture donor was not significant.

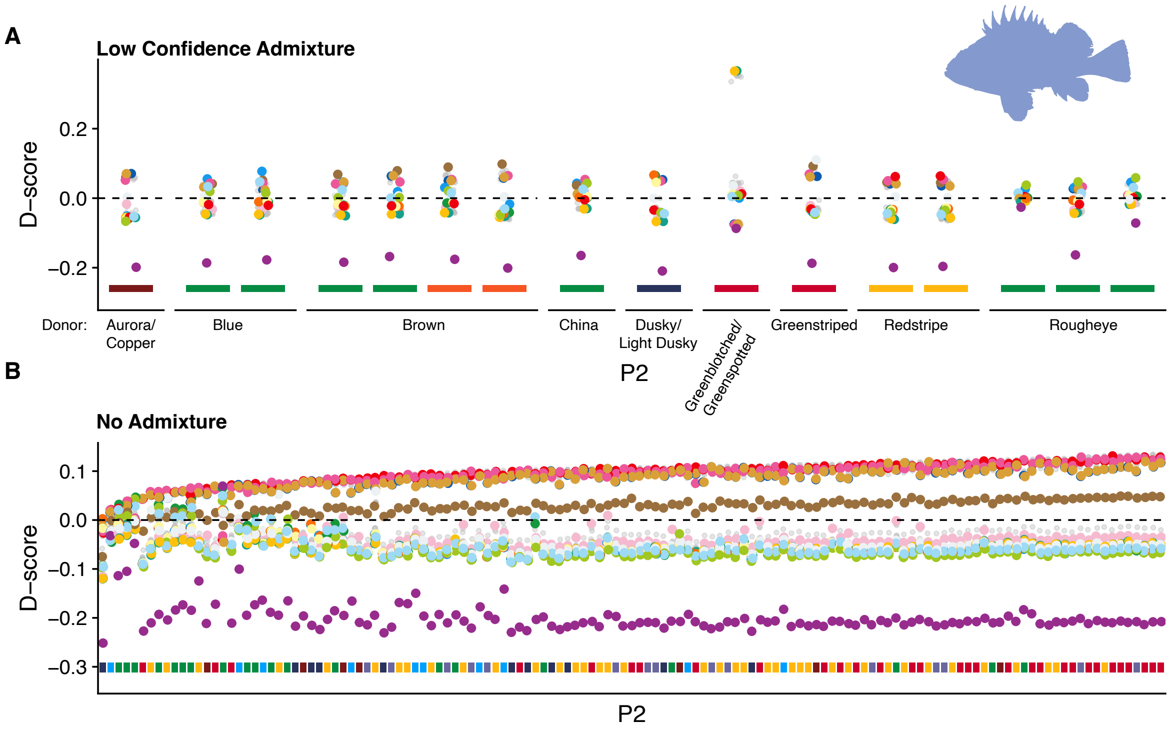


**Figure S10: Introgression into Quillback from diverse sources.** D scores for Quillback samples with all species as the possible donor. Each point represents the trio (All other Copper, Tested Copper, Donor species). The bar under each sample represents its geographic location (See Figure 2). Species not identified as being involved in admixture are coded are smaller and grey for clarity. A) Low confidence admixture: where the gap between the 1st and 2nd ranked D-score is between 0.01 and 0.02. B) No admixture: where the gap between the 1st and 2nd ranked D-score is less than 0.01, or the top admixture donor was not significant.
